# EGFR-targeted affibody–polyIC polyplex kills EGFR-overexpressing cancer cells without activating the EGFR

**DOI:** 10.1101/2025.10.01.679886

**Authors:** Anne Pettikiriarachchi, Yelena Ugolev, Richard Birkinshaw, Ahmad Wardak, Maree C. Faux, Timothy E. Adams, Salim Joubran, Alexei Shir, Nufar Edinger, Maya Zigler, Antony W Burgess, Alexander Levitzki

## Abstract

The epidermal growth factor receptor (EGFR) is aberrantly activated in many human epithelial cancers. This report presents the preparation, purification, and the anti-cancer potency of an anti-EGFR affibody (Z_EGFR 1907’_)-polyethylenimine (PEI)-polyIC complex (PPEA-polyplex). Surface plasmon resonance analysis showed that the Z_EGFR 1907’_ affibody binds tightly to full-length sEGFR with an average equilibrium dissociation constant, K_D,_ value of 6.74 nM. The PPEA-polyplex does not activate the EGFR kinase, but kills tumor cells expressing medium to high levels of EGFR. The PPEA-polyplex stimulates the release of chemotactic cytokines (e.g., GRO-α, IFN-γ–inducible protein-10) and promoted PBMC-mediated bystander killing of non-transfected tumor cells. The PPEA-polyplex inhibited the growth of human epidermoid vulval carcinoma (A431) xenografts growing in immunocompromised nude mice. PPEA-polyplexes have the potential to inhibit the growth of tumors in Triple-negative breast cancer (TNBC) patients and other cancers which over-express the EGFR.

## Introduction

The epidermal growth factor receptor (EGFR) was identified as a target for cancer therapy almost forty years ago(1). Over 70% of malignant tumors have abnormal expression or activation of the EGFR, including brain(2), bladder(3), colorectal(4, 5), head and neck(6), lung(7) and breast cancers(8). Cancers can be driven by autocrine activation of the receptor(9, 10), mutation(11) or amplification(12) of the EGFR gene. This perturbation of EGFR signaling homeostasis plays a critical role in initiation, progression, growth and metastasis of most colorectal cancers (CRCs)(13, 14). Targeting cancers with EGFR receptor antagonists (e.g. antibodies or specific kinase inhibitors) has already helped millions of cancer patients world-wide(15). In particular CRC patients without K-ras mutations and lung cancer patients with mutant EGFR have responded to EGFR antagonist treatment(16). There are indications that combined, targeted therapies focused on the EGFR family might help many other cancer patients(17), including pancreatic(18), head and neck(19) and brain cancer patients(20).

Cancer immunotherapy has progressed, especially through the use of anti-CTL4A and anti-PD1/PDL1 in the treatment of metastatic melanoma and the emerging success of tumor targeting chimeric T-cells(21, 22). We have previously demonstrated that EGFR-targeted dsRNA polyinosine/polycytosine (polyIC) eradicates EGFR overexpressing tumors in experimental animals(23, 24). PolyIC has been known as an immune stimulating agent for dozens of years but was too toxic for systemic utilization. Yet, polyIC is used by local application to enhance the action of vaccines(25, 26).

Tumor cells expressing high levels of EGFR can be killed with EGFR-homing chemical vectors loaded with polyIC(23, 27). In these previous studies(23, 24, 27) EGF was used to deliver the polyIC to the surface of the neoplastic cells and following endocytosis the delivery of the polyIC to the cytoplasm was facilitated by the polyethyleneimine-polyethylene-glycol (PEI-PEG) linker (24). This PEI-PEG-EGF (PPE) polyIC complex (polyplex) induced tumor cell killing, both *in vitro* and *in vivo*. Moreover, we showed that following internalization, polyIC activates multiple cell-killing mechanisms and induces strong bystander effects, leading to killing of both targeted and non-targeted tumor cells, without harming the neighboring normal cells(23, 24, 27). The resistance of normal cells, including cells which express low numbers of EGFR, is most likely due to the more robust nature of these cells as compared to tumor cells, which are under constant stress(28).

In 2007 Friedman and colleagues engineered a series of EGFR-binding affibodies(29): Z_EGFR 1907_ was determined to bind to the EGFR with high affinity, K_D_ of 5.4 nM. Optimisation of the affibody scaffoldings led to development of a protein with improved properties(30) and further modifications resulted in an improved variant of Z_EGFR 1907_ with increased thermal stability, Z_EGFR 1907’_. The Z_EGFR 1907’_ affibody binding to EGFR does not appear to induce receptor phosphorylation, but it is internalised by the cell(31, 32). In this report we engineer, characterise and evaluate the potency of Z_EGFR 1907’_-polyIC-polyplexes for targeting the EGFR. The Z_EGFR1907’_ affibody was linked covalently to a LPEI-PEG diconjugate to form the LPEI-PEG-Z_EGFR1907’_ triconjugate which targets the EGFR. The Z_EGFR 1907’-_affibody-triconjugate (PPEA) was complexed with polyIC to form the anti-tumor Z_EGFR 1907’_-affibody-polyIC complex (PPEA-polyplex). The potency of the anti-tumor activity of the PPEA-polyplex was investigated.

## Abbreviations

Abbreviations are appended to the full description of the chemicals when they first appear in the text, however it is important to note that the abbreviation PPEA-polyplex is a conjugate of polyethyleneglycol, polyethyleneimine, polyIC and the anti-EGFR affibody Z_EGFR1907’_. When it is appropriate to emphasize that this polyplex contains polyIC, we also use the term PPEA-polyIC-polyplex for this conjugate. Z_EGFR 1907’_ is used as an abbreviation for Z_EGFR 1907’_ affibody.

## Materials and Methods

Plasmid DNA encoding the affibody Z_EGFR 1907’_-Cys (an affibody which binds to the EGFR) was transformed into *Escherichia coli* BL21 (DE3). The transformed bacteria were grown overnight in 50 ml 2xYT media with 1% glucose and 30 µg/ml kanamycin at 37°C. This overnight starter culture was transferred into larger culture and allowed to grow to an OD_600nm_ between 0.5 and 1.0. Protein expression was induced with 0.5 mM isopropyl-β-D-1-thiogalactopyranoside (IPTG) - overnight at 28°C. The bacterial cell pellet was stored at -80°C until required for the purification processes.

The bacterial cell pellet containing the anti-EGFR affibody was thawed and resuspended in 50 ml 20 mM Hepes pH 7.4, 500 mM NaCl, 10% glycerol, 10 mM Imidazole and 2 mM β-mercaptoethanol (Buffer A) and lysed using a homogeniser. The soluble proteins were recovered by centrifugation at 20,000 rpm for 20 min at 4°C. His-Tag EGFR affibody protein was purified using a Ni-NTA column. The column with the bound anti-EGFR affibody was washed with Buffer A for 15 column volumes (CV) and with Buffer A containing 30 mM Imidazole for 10 CV. Bound EGFR affibody was eluted using buffer A containing 500 mM Imidazole for 5 CV. The purity of the EGFR Affibody protein was assessed by SDS-PAGE analysis (Figure 1Ai). The eluted protein was then loaded onto a gel filtration chromatography column Superdex 75 16/60 (GE healthcare) equilibrated using 20 mM Hepes pH7.4, 500 mM NaCl, 10% Glycerol and 2 mM β-mercaptoethanol. 1 ml fractions from the peak spectrum were collected and the purity of the gel filtration purified samples were assessed by SDS-PAGE (Figure 1Bi and1Bii). Ellman’s reagent (Thermo Scientific) was used to determine the presence of a sulphydryl group as per the manufacturer’s protocol.

**Figure 1.**
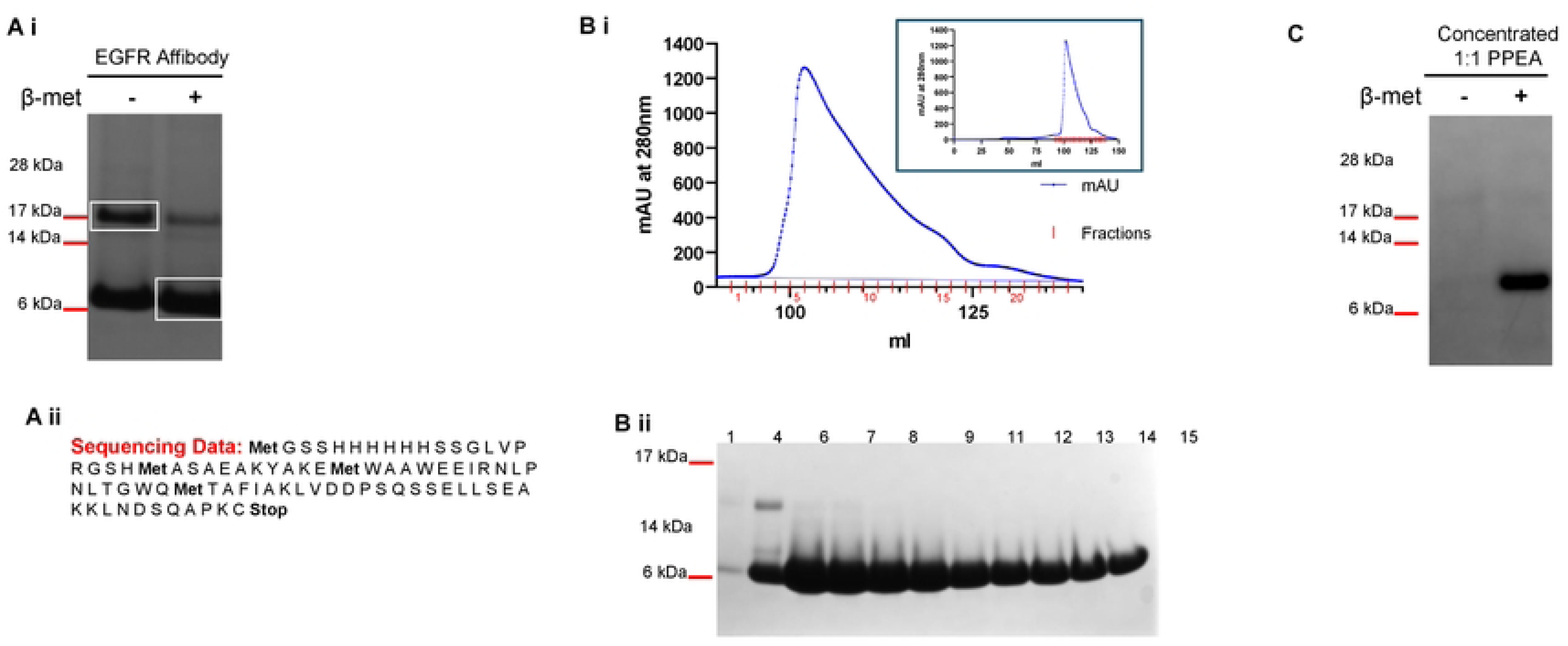
Generation of the LPEI-PEG-Z_EGFR1907’_. (**A i**) SDS-PAGE analysis showing the purity of the Ni-NTA purified Z_EGFR 1907’_ under pseudo native and denatured conditions. Pseudo native gel bands (boxed) were extracted and analysed by mass spectrometry. Mass spectrometry data confirmed that both bands correspond to one species, (**A ii**) DNA sequencing of the plasmid revealed the above sequence and this was also confirmed by mass spectrometry analysis (**B i**) Elution profile of the size exclusion gel filtration chromatography of Z_EGFR 1907’_ affibody purified via His-Tag Ni-NTA purification. Superdex 75 1660 column equilibrated with 20 mM Hepes pH 7.4, 500 mM NaCl, 10% Glycerol and 2 mM β mercaptoethanol and 1 ml fractions were collected. Insert shows the complete elution profile. (**B** ii) 20 µl of selected sample fractions from the purification were denatured, run on an SDS-PAGE gel and stained with coomassie blue. (**C**) Silver stained SDS-PAGE gel of the concentrated 1:1 triconjugateZ_EGFR 1907’_ (PPEA) under pseudo native and denatured conditions. Pseudo native and denatured conditions are indicated by -/+ β-mercaptoethanol.

### SDS-polyacrylamide gel electrophoresis (SDS-PAGE) analysis of proteins

Samples of Ni-NTA–purified Z_EGFR-1907’_ were diluted in SDS loading buffer with or without 1.43 M β-mercaptoethanol, heated to denature, and separated on NuPAGE 4–12% Bis-Tris gels (MES buffer; Invitrogen). Protein bands were stained with Coomassie Brilliant Blue R-250 (BioRad) (Figure 1Ai). The concentrated 1:1 Triconjugate Z_EGFR-1907’_ affibody construct (see Synthesis of the LPEI-PEG-EGFR Affibody section below) was also analysed using SDS-PAGE. To visualise the sample bands, the gel was silver stained as follows: washed with 40 mM DTT, stained with 1 mg/ml AgNO_3_, washed with MilliQ, developed with 30 mg/ml Na_2_CO_3_ and 0.15% formaldehyde, reaction stopped with MilliQ and gel fixed with 10% acetic acid (Figure 1C).

### Mass Spectrometry Analysis

Protein bands responding to the Z_EGFR 1907’_-affibody were extracted from a Coomassie stained SDS-PAGE gel and subjected to manual in-gel tryptic digestion. Briefly, the gel spots were de-stained and dehydrated with acetonitrile before being submitted to reduction with 50 mM TCEP solution (50 μL of 0.5 M TCEP / 125 μL of 200 mM TEAB / 325 μL milliQ water) and alkylation with 100 mM Iodoacetamide solution in 50 mM TEAB buffer. Trypsin solution was prepared by adding 50 μL of 50 mM TEAB to each trypsin single vial (1 μg trypsin) and rehydrating dried gel plug/bands with approximately 10 μL trypsin working solution for 60 minutes at 4°C. Excess trypsin solution was removed with a pipette before adding 35 μL of 50 mM TEAB to the sample vial to ensure proper hydration during digestion at elevated temperatures. Digestion was performed at 37°C overnight. The digestion was stopped by adding 5 μL of 10% formic acid before collecting the supernatant. The generated tryptic peptides were concentrated to ∼20 μL by centrifugal lyophilization for LC MS/MS analysis for peptide mapping mass spectrometry on LC/MS/MS.

Mass spectrometry was used to confirm protein purity, molecular mass and the protein sequence (Figure 1Aii). A spectrometer (Thermo Scientific) with a nanoESI interface in conjunction with an Ultimate 3000 RSLC nano HPLC (Dionex Ultimate 3000) was used. The LC system was equipped with an Acclaim Pepmap nanotrap column (Dionex-C18, 100 Å, 75 μm x 2 cm) and an Acclaim Pepmap RSLC analytical column (Dionex-C18, 100 Å, 75 μm x 15 cm). The tryptic peptides were injected into the enrichment column at an isocratic flow of 5 μL/min of 3% v/v CH3CN containing 0.1% v/v formic acid for 5 min applied before the enrichment column was switched in-line with the analytical column. The eluents were 0.1% v/v formic acid (solvent A) and 100% v/v CH3CN in 0.l% v/v formic acid (solvent B). The flow gradient was (i) 0-5 min at 3% B, (ii) 5-6 min, 3-6% B (iii) 6-18 min, 6-10% B (iv) 18-38 min, 10-30% B (v) 38-40 min, 30-45% B (vi) 40-42 min, 45-80% B (vii) 42-45 min at 80% B (viii) 45-46 min, 80-3% B and (ix) 46-53 min at 3% B. The LTQ Orbitrap Elite spectrometer was operated in the data-dependent mode with nano ESI spray voltage of 2.0 kV, capillary temperature of 250°C and S-lens RF value of 55%. All spectra were acquired in positive mode with full scan MS spectra scanning from m/z 300-1650 in the FT mode at 240,000 resolutions after accumulating to a target value of 1.0x10^6^. Lock mass of 445.120025 was used.

The top 20 most intense precursors were subjected to collision-induced dissociation (CID) with normalized collision energy of 30 and activation. The generated MGF files were uploaded to the Bio21 MSPF Pipeline (Bio21 Mass Spectrometry and Proteomics Facility Pipeline, http://proteomics.bio21.unimelb.edu.au/msile) and then searched using the MASCOT version 2.4.01 for algorithm against the murine Uniprot fasta database that included the added affibody fasta sequences (84562 sequences in total). The search parameters consisted of carbamidomethyl of cysteine as a fixed modification, NH2-terminal acetylation and oxidation of methionine as variable modifications. Up to three missed tryptic cleavage sites were allowed (trypsin/P) with a peptide mass tolerance of 20 ppm, a fragment mass tolerance of 0.8 Da, automatic decoy searches enabled with low decoy rate (< 5%), a significance threshold p < 0.05 and a Mascot score >30.

### Biosensor Analyses

SPR binding assays (Biacore S200, Cytiva) were performed to measure interactions of Z_EGFR 1907’_and cetuximab withsEGFR_1-501_ and sEGFR_1-621_. Proteins were immobilized on CM5 chips by amine coupling (pH 5.0) to ∼1500 RU (sEGFR_1-501_) and ∼1700 RU (sEGFR_1-621_).An initial pH scouting experiment was performed using 10 mM Acetate with pH 4.0, 4.5, 5.0, 5.5 and 6.0 for sEGFR proteins, sEGFR_1-501_ and sEGFR_1-621_. Samples were injected over the flow cell surface at 10 µL/min for 12 min. EGFR proteins were diluted in SPR buffer, 10 mM Hepes pH 7.4, 150 mM NaCl, 3 mM EDTA and 0.005%(v/v) Tween 20. EGFR proteins, sEGFR_1-501_ and EGFR_1-621_, were immobilized at 1500 RU and 1700 RU (resonance units) respectively to Series S Sensor Chip CM5 (Cytiva) by amine coupling at pH 5.0 (Table S1), according to the manufacturer’s instructions.

The conditioning cycle consisted of 0.1% SDS for 60 sec, 100 mM HCl for 60 sec, 10 mM NaOH for 60 sec and 50 mM glycine pH 2.5 for 60 sec. For binding kinetic studies, proteins were injected at 30 µL/min for 300 sec association over the sEGFR surfaces. Z_EGFR 1907’_ and Cetuximab concentrations ranged from 0 to 250 nM. This was followed by 800 sec dissociation phase. After each injection, the sensor surface was regenerated with SPR buffer supplemented with 50 mM glycine pH 2.5 for 120 sec between each cycle before repeating the sample injection. The dissociation equilibrium constant (K_D_) was calculated using Biacore S200 Evaluation Software 1.1 (Cytiva). The curves were fitted using a local 1:1 binding model and a global heterogenous binding model.

The binding sites of the Z_EGFR 1907’_ and Cetuximab to sEGFR_1_-_621_ were explored. For A-B-A competitive binding assays, various concentrations and combinations of competitor sample solutions (B from A-B-A assay) and flanking competitor sample solutions (A from A-B-A assay) were tested. The assay consisted of 200 sec of flanking solution A, followed by 180 sec of sample solution B and further 60 sec of flanking solution A. Protein samples were pumped over the cell at a rate of 30 µL/min. The sensorgrams generated were subtracted from reference flow cell values. After each condition the flow cell was regenerated with SPR buffer supplemented with 50 mM glycine pH 2.5.

### Synthesis of LPEI-PEG-OPSS thiol reactive co-polymer

Linear polyethyleneimine (LPEI) precursor, poly(2-ethyl-2-oxazoline), was synthesized following the methods published by Brissault and colleagues(33, 34). In brief, methyl p-toluenesulfonate (74 mg, 397 mmol) dissolved in 60 ml acetonitrile was added to 8 ml (79 mmol) of 2-ethyl-2-oxazoline. The reaction mixture was stirred under reflux for 2 hours, during which time a precipitate formed. The crude product was dissolved in 100 ml of methylene chloride and precipitated in 400 ml of diethyl ether yielding 5.5 g of poly(2-ethyl-2-oxazoline) as a yellow powder.

Synthesis of the LPEI (free base form) was prepared as described previously(34, 35). In brief, 5.5 g (0.11 mmol) of poly(2-thyl-2-oxazoline) was hydrolyzed with 68.8 ml of concentrated HCl (37%). The resulting air-dried LPEI hydrochloride salt was dissolved in 100 ml of water to yield 4 g of LPEI salt. This was made alkaline by adding 100 ml of 3 M NaOH. The lyophilised solid weighted ∼1.25 g. LPEI-PEG_2k_-OPSS diconjugates (1:1 and 1:3) were synthesized as previously described(35). 174 mg (8 µmol) of LPEI in 2.7 ml of absolute EtOH was mixed with 79 mg of OPSS-PEG2k-CONHS (39.5 µmol) in 500 µl of anhydrous DMSO. The co-polymers conjugated with different molar ratios of PEG to LPEI were separated by cation exchange chromatography using a HR10/10 column packed with MacroPrep High S resin (BioRed).

Upon agitation for 3 hrs, 2 ml of 20 mM Hepes pH 7.4 were added to the reaction mixture and after few minutes, once the viscosity increased, the pH was adjusted to 7.4 with 1 M HCl. The column was equilibrated with 20 mM Hepes pH 7.4 buffer (buffer A). Separation was performed under a 20 mM Hepes pH 7.4; 3 M NaCl (buffer B) gradient. Sample was loaded at 1 ml/min for 7 min and for further 60 min with 0% buffer B. This was followed by isocratic elution at 20% buffer B until 76 min, followed by gradient elution up to 45% buffer B at 87 min and continuing at 45% buffer B until 110 min. Then, isocratic elution at 50% buffer B until 120 min was followed by gradual increase in gradient up to 100% buffer B until 180 min. Isocratic elution continued at 100% buffer B until 215 min, followed by a gradual decrease in gradient to 0% buffer B by 218 min. 2 ml fractions were collected between 0% -50% Buffer A and 1 ml fractions were collected during the elution step (50% -100% buffer B). Sample separations were monitored at 220 nm, 280 nm and 343 nm, and all fractions from the three peaks were stored -20°C.

Fractions from the 1:1 PEG to LPEI conjugation, which eluted during the 50% to 100% buffer B, were combined and desalted against 20 mM Hepes buffer pH 7.4 using a 20 ml HiTrap desalting column (4 x 5 ml Sephadex G25 columns). The purest fractions containing LPEI-free LPEI-PEG-OPSS were pooled and stored in the dark at -20°C. A copper assay(36) was used to determine the concentration of di-conjugate 1:1, with molar ration of LPEI to PEG ∼1:1. In brief, copper assay consisted of incubating copolymers (initially dissolved at 2x concentration with 100% methanol and then diluted to 1x with MilliQ) with CuSO_4_ (23 mg of CuSO_4_.5H_2_O in 100 ml of 0.1 M acetate buffer pH 5.4) for 20 min and measuring their absorbance at 285 nm.

### Synthesis of the LPEI-PEG-EGFR Affibody

The 1:1 diconjugate was mixed with EGFR affibody as described by Joubran *et al*(35). The reaction mixture was vortexed at ∼800 rpm for 24 hrs and stored at -20°C. Later, the reaction mixture was thawed and purified using cation exchange chromatography using a HR10/10 filled with MacroPrep High S resin (BioRad). 3 step gradient elution from 20 mM Hepes pH 7.4 reaching to 20 mM Hepes pH 7.4 with 3 M NaCl was used. Sample was loaded at 1 ml/min for 6 min and for further 10 min with 0% buffer B. Then isocratic elution with 33% buffer B until 15 min was followed by gradient elution reaching 66% buffer B at 30 min, isocratic elution from 30 min to 37 min, and gradient elution reaching 100% buffer B at 55 min. Finally, there was a gradual decrease down to 33% buffer B at 60 min. 1 ml fractions were collected between 0% -100% Buffer B. Sample separation was monitored at 213 nm, 280 nm and 343 nm, and all fractions from the three peaks were stored at -20°C. The 1:1 triconjugate fractions eluting between 25 min and 32 min (i.e. the last peak eluted by the gradient) were combined and desalted using the 4x5 ml Sephadex G25 (two runs) against PBS. Desalted fractions were combined, and 1 ml aliquots were frozen at -80°C. The concentration of the 1:1 triconjugate was determined using the copper assay. The triconjugate fractions were pooled, concentrated and buffer exchanged to 20 mM Hepes pH 7.4, 5% glucose with RNase free MilliQ (HBG) using a 3K MWCO Amicon Ultra15 Centrifugal filter unit. The presence of the affibody (∼8 kDa) was confirmed under denaturing/reducing conditions (Figure 1C).

### EGFR Affibody polyIC-Polyplex Formation

The 1:1 triconjugate affibody was complexed with poly (I:C) (Tocris, UK) in HBG buffer at a Nitrogen to Phosphate (N/P) ratio of 6 (where N=nitrogen from LPEI and P= phosphate from polyIC (35)). This ratio corresponds to a LPEI/polyIC weight ratio of 0.78(37). Both the 1:1 Triconjugate Affibody and polyIC were diluted up to twice the final volume using HBG buffer. Fresh polyplexes were made prior to the experiment by transferring equal volumes of 1:1 Triconjugate to the diluted polyIC and mixing by pipetting. The polyplexes were incubated at room temperature for 30 mins. The final 1:1 triconjugate Affibody-polyIC-polyplex (Z_EGFR 1907’-_polyIC-polyplex) was determined to be 100 µg/ml using the copper assay [36].

### Polyplex Size Measurements

To determine the influence polyIC concentration has on the polyplexes formed, the particle size and its distribution were measured at 25°C by dynamic light scattering using a Zetasizer µV (Malvern Instruments). The instrument was fixed with an 830 nm laser wavelength and the optic arrangement angle at which the measurements were performed was 90°. Each polyplex sample was measured in triplicate.

### Cell Culture

EGFR-overexpressing A431 human epidermoid vulval carcinoma (38), SK-BR-3(39), U87MG human glioma and its EGFR-overexpressing subline U87MGwtEGFR (40) cell lines were grown in Dulbecco’s modified Eagle’s medium (DMEM). U87MGwtEGFR cells were supplemented with geneticin sulfate (G418, Gibco, UK) at a final geneticin concentration of 500 µg/ml. MCF7, MDA-MB-231, MDA-MB-468 and BT-474(41) cells were grown in Roswell Park Memorial Institute (RPMI). U138MG cells were grown in Eagle’s Minimum Essential Medium (MEM) supplemented with 2 mM L-glutamine, non-essential amino acids and 0.1 g/L sodium pyruvate. All cell culture media were supplemented with 10% fetal bovine serum (FBS),100 U/mL penicillin, and 100 µg/mL streptomycin and all cells were grown at 37 °C in an incubator with humidified air equilibrated with 5% CO_2_.

CRC cell lines: Difi, SW620, Lim2537, SW480, Lim2099, HCA7, Colo201, Colo320, HCT116, IS1, SW48, MC38, SW1116 and CX1 were obtained from colleagues either at WEHI or the Olivia Newton John Cancer Research Institute (ONJCRI). Other cell lines such as A431, were obtained from colleagues at WEHI, BT-20 and MDA-MB-468 were purchased from ATCC. All cell lines were grown in their recommended media (DMEM from Gibco, RPMI 1640 from Sigma and EMEM from ATCC) supplemented with 10%FCS, -/+ Adds (1.08% thioglycerol, 50 mg/ml hydrocortisone, 100 U/ml insulin), -/+ G418. All cell lines were grown to approximately 75% confluency before passaging. All cell lines except MC38, BT-20 and MDA-MB-468 cell lines were maintained at 37°C and 10% CO_2_. MC38, BT-20 and MDA-MB-468 cell lines were grown at 37 °C and 5% CO_2_. The original sources, current locations of these CRC cell lines and the characteristics of each cell line are documented in the Supplementary material of Wang *et al.*(42).

### Cell Viability Assay

Briefly, the optimal seeding cell density for each cell line was determined in triplicate. The cells were treated the same way as for the CellTiter Glo assay without drug (see below).

After treatment with the PPEA-polyplexes, cell proliferation was determined using a CellTiter-Glo assay (Promega, Madison) on a range of colorectal cancer (CRC) cell lines, A431, BT-20, MDA-MB-468 and MC38 sub-clones. A cytotoxic drug, 1 µM Bortezomib or WEHI-7326(43), was used as the total death control and the untreated cell viability was determined by adding an equivalent volume of the HBG buffer. For the CellTitre Glo assay, cells were plated in triplicates in 96 well plates and after 24 hr were treated with drugs for 72 hrs. Cell survival was quantitated using 2D CellTiter-Glo assay (Promega) and luminescence plate reader (EnVision 2100 Multilable reader) with Wallac EnVision Manager 1.12 software.

### FACS-based EGF receptor quantitation

For FACS analysis 0.5 x 10^6^ cells in 50 µl were seeded into U-bottom 96 well plates and the cell suspension was treated with 2.5 µg/ml mouse anti-human EGFR monoclonal antibody 528 (WEHI) and 0.4 µg/ml F(ab’)2-Goat anti-Mouse IgG (H+L), PE, (eBioscience) or 1.5 µl of PE anti-human EGFR antibody (BioLegend). Cells were incubated on ice for 1hr for anti-mouse 528 and 30 mins for PE antibody or 45 min for PE anti-human EGFR antibody. All samples except control cell preparations were stained with either 5 µg/ml Propidium Iodide (Thermofisher) or 0.1 µg/ml DAPI (Thermofisher). Cells expressing fluorescent protein and/or cell surface markers were analysed using CytoFLEX (Beckmann Coulter) or Fortessa1 (BD Biosciences) flow cytometers. Each analysis consisted of at least 10,000 live cells (events). Spectral bleed-through between the PI channel and PE channel were corrected by compensation. A431 and Difi cells were run as positive controls every time. FlowJo 10 software (BD Biosciences) was used for FACS data analyses. Cells were gated in order: live cells, single cells and PI or DAPI negative cells. The final sorted population (PI negative cells) was analysed for presence of GFP using the 525 nm channel and PE using the 585 nm channel, and calculations on fluorescence intensity performed. The median values for the 585 nm population in quadrant 3 were obtained by subtracting the median PE value from median 528 nm, i.e.1°AB + PE. The level of EGFR expression was expressed as a percentage of A431 or Difi EGFR expression. The error bars represent the robust standard deviation of the difference between sample medians (σ_d_) (44). σ_d_ was calculated using the formula = sqrt (σ_1_ ^2^ / n + σ ^2^ / n) where σ and σ refers to the robust standard deviations from sample 1 (528 1°AB + PE) and sample 2 (PE). n_1_ and n_2_ refers to the number of cells measured in each of these samples.

### PBMC Extraction

Peripheral blood mononuclear cells (PBMC) from healthy human donors were separated on Ficoll Plaque (Pharmacia). Briefly, cells were diluted in RPMI 1640 (x4) and 40 mL of diluted cell suspension was layered over 13 mL of Ficoll Plaque in 50 mL conical tubes. Following centrifugation at 400g for 30 mins at 20 °C in a swinging bucket rotor without brakes, the mononuclear cell layer was transferred to a new 50 mL conical tube, washed twice with 50 ml RPMI 1640 medium, resuspended at a density of 4x10^6^ cells/ml and cultured in this RPMI 1640 medium supplemented with 10% fetal calf serum, 100 units/ml penicillin and 100 µg/ml streptomycin.

### Chemotaxis Assay

Chemotaxis was performed as described previously(27), except that chemotaxis plates were from Corning (HTS Transwell-96 well plate, 5 µm polycarbonate membrane, catalogue no.3388).

### Cytokine Measurements by ELISA

Supernatants from polyIC-treated cells were harvested and assayed for IP-10 and Gro-α using commercial ELISA kits (PeproTech) according to the manufacturer’s instructions.

### Cell Treatment and Immunoblotting

MDA-MB-468 cells were seeded in 6-well plates (600,000 cells/well), allowed to grow for 24 hours in compete medium and starved for 24 hours in serum-free medium. PPEA was diluted in HBS (20 mM Hepes pH 7.4, 150 mM NaCl) and cells were treated in duplicates with 120 ng PPEA for different periods of time. As a positive control for EGFR kinase activity, cells were treated with 10 ng EGF for 5 min. Cells were lysed with boiling sample buffer (10% glycerol, 50 mmol/L Tris-HCl pH 6.8, 3% SDS, and 5% 2-mercaptoethanol). Protein determination was performed using the bound Coomassie Blue method, as described previously(45). Lysates containing 60 µg of protein were resolved on 10% SDS-PAGE gel. Western blot analysis was conducted as described previously(46). The following antibodies were used: Antibody specific for EGFR was purchased from Santa Cruz Biotechnology (catalogue no.sc-03, dilution 1:1000); Antibody recognizing phosphorylated EGFR (Tyr1068) was from Cell Signaling Technology (catalogue no. 2234S, dilution 1:1000); Anti-GAPDH was from Cell Signaling Technology (catalogue no. 2118, dilution 1:1000).

### Treatment of Subcutaneous Xenografts & Allografts

Female nude mice (NUDE-HSD: Athymic Nude-NU mice) aged 3-4 weeks were obtained from Harlan, Rehovot. All animal experiments were performed according to the Hebrew University Ethical Committee regulations.

For A431 xenografts, two million A431 cells resuspended in 0.2 ml of PBS were injected subcutaneously into the right flank of immune compromised female athymic nude mice. The volume of growing tumors was calculated as follows: V=LW^2^/2 (L=length and W=width). When the tumors reached an average volume of 137 mm^3^, mice were randomly divided into 5 groups (n=7 or 8 per group), and the treatment was initiated. PolyIC formulated with PPEA in HBG buffer at w/w ratio of 0.78, which correlates with N/P ratio (molar ratio of nitrogen in PEI to phosphate in polyIC) of 6 as described previously(23, 27), was injected. The polyplexes were administered via intravenous injection at doses of 0.75 mg/kg or 0.1 mg/kg every 24 hours, 6 days per week, for a total of 9 injections over a 10-day period. Tumor volumes were measured on days 3, 6 and 10 after starting the intravenous injections.

## Results

### Engineering and characterisation of triconjugate affibody-polyIC-polyplexes

We engineered an affibody-PEG-polyIC-polyplex to target the EGFR and deliver polyIC to the cytoplasm of cancer cells. An analogue of the anti-EGFR affibody(29), Z_EGFR 1907’_-Cys was expressed in *E.coli* BL21 (DE3) and purified on a Ni-NTA column from lysates as described in the methods section. SDS-PAGE analysis of the sample detected two bands with apparent molecular weights ∼8 kDa and 16 kDa under pseudo native conditions (Figure 1A; see Materials and Methods). These bands were analysed by mass spectrometry and the resulting sequence for the Z_EGFR 1907’_-Cys affibody is shown in Figure 1A. The 8 kDa and 16 kDa bands correspond to the monomer and dimer of the Z_EGFR 1907’_-affibody. The monomeric form of the Z_EGFR 1907’_-Cys affibody was purified by gel filtration (Figure 1B). All subsequent experiments were performed using the purified with the Z_EGFR 1907’_-Cys affibody monomer.

### Binding of the Z_EGFR 1907’_-Cys affibody to the extracellular domain of the EGFR

Surface plasmon resonance analysis was used to measure the binding of the monomeric Z_EGFR 1907’_-Cys affibody to two forms of the extracellular domain of EGFR: sEGFR_1-501_ (a truncated form of the extracellular domain)(47) and sEGFR_1-621_ (the full length extracellular domain)(48–50) (Figure 2A). Our binding kinetics data indicates that Z_EGFR 1907’_-Cys affibody binds more tightly to EGFR_1-621_ than EGFR_1-501_ and has an average K_D_ value of 6.74 ± 0.91 nM SD (Supplementary Figure 2A and Table 1). Z_EGFR 1907_ has been reported previously to have a K_D_ value of 5.4 nM using a similar method(29), however, different EGFR analogues were immobilized onto the biosensor. The strong affinity of the Z_EGFR 1907’_-Cys affibody to EGFR_1-621_ is consistent with having a more accessible groove created by folding of domain I and III in an untethered confirmation by the full-length extracellular domain sEGFR_1-621_ (51). However, fitting the data to a global model suggests that the Z_EGFR 1907’_-Cys affibody binds to two sites on the sEGFR_1-621_ receptor with two different binding affinities (Table S3 and Supplementary Figure S1 and S2C). This was not evident when affibody bound to sEGFR_1-501_. This finding supports the notion that sEGFR_1-621_ adopts two conformations, tethered and untethered(52–54).

**Figure 2.**
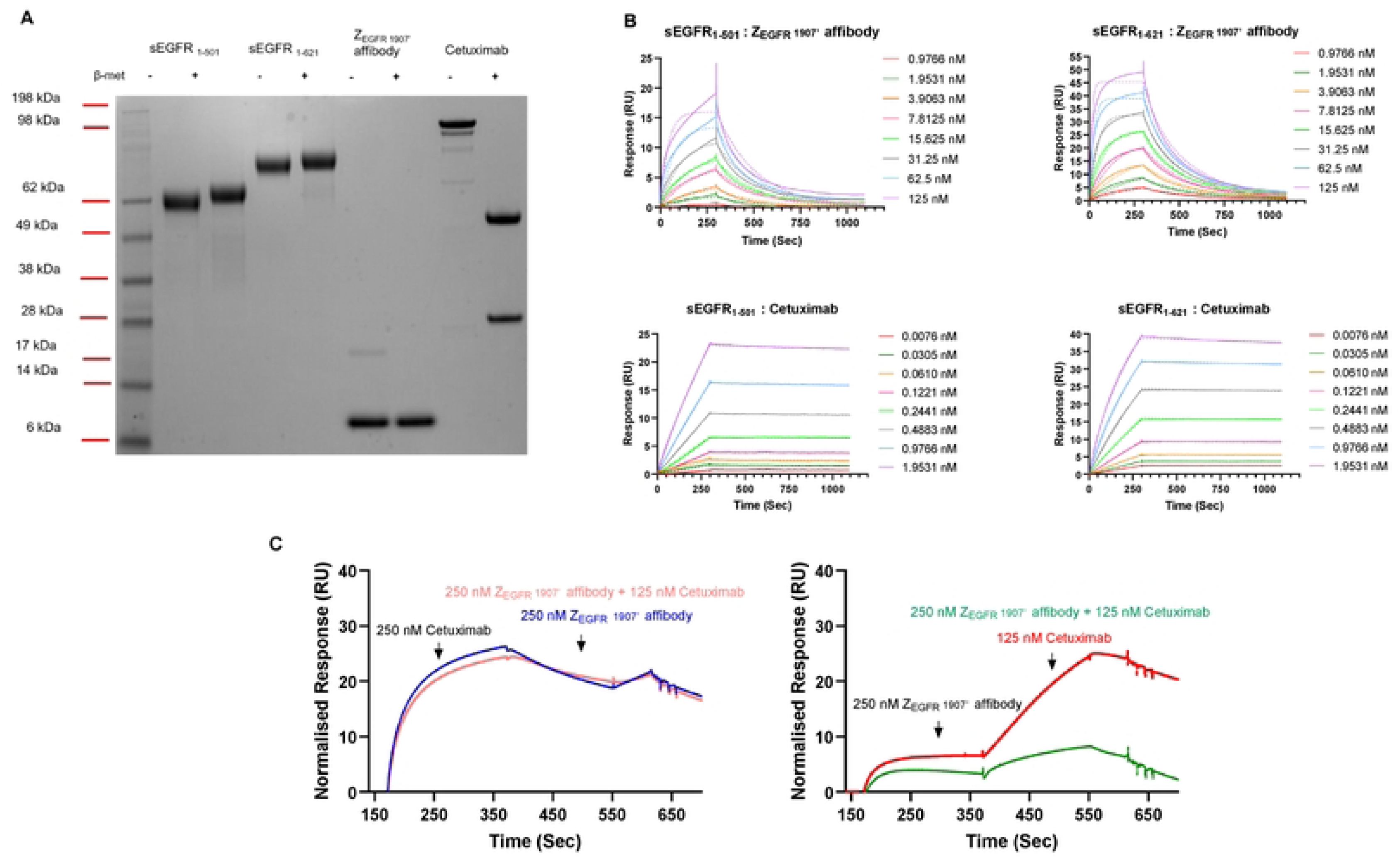
Biacore analysis of Z_EGFR 1907’_binding to EGFR proteins. SDS-PAGE analysis showing the purity of the proteins used for the Biacore analyses. Expected molecular weights: sEGFR_1-501_ 56kDA, sEGFR_1-621_ 69kDa, Z_EGFR 1907’_11kDa and Cetuximab 152kDa. **(B)** Binding kinetics for the Z_EGFR 1907’_ and Cetuximab interacting with sEGFR_1-501_ andsEGFR_1-621_, used to determine K_d_ values using Biacore S200.**(C)** Competitive binding kinetics of Z_EGFR 1907’_ and Cetuximab detected under various conditions shows Cetuximab andZ_EGFR 1907’_ appear to bind to the same EGFR binding site. Solid and dotted line graphs represent raw and fitted data respectively.

**Table 1.**
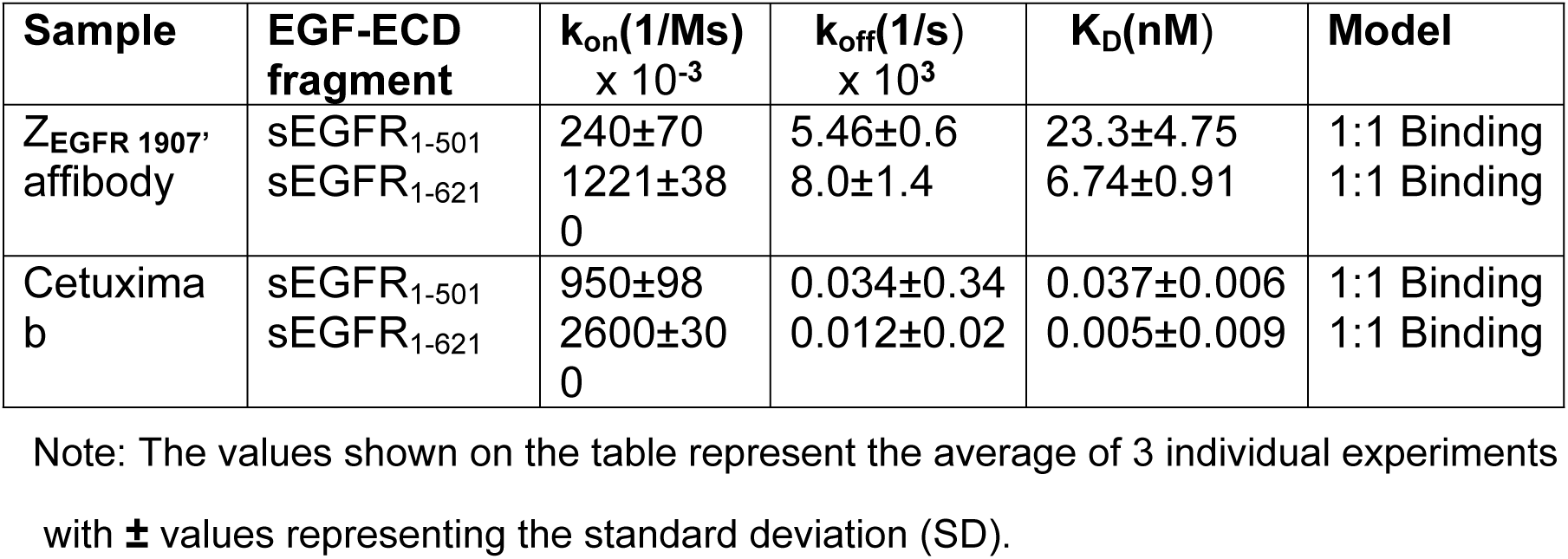
Binding affinities of Cetuximab and Z_EGFR 1907’_ affibody to the EGFR-extracellular domain (ECD).

### Comparison of the binding of the ZEGFR 1907’ and Cetuximab to sEGFR

For comparison purposes we studied the binding kinetics of Cetuximab(55, 56) binding to the EGFR extracellular domain. Binding kinetic curves for Cetuximab showed that once bound to either sEGFR proteins, Cetuximab is slow in coming off the receptor as indicated by the flat dissociation curves (Supplementary Figure 2C and S2B). Comparison between ZEGFR 1907’ and Cetuximab shows Cetuximab does not come off the EGFR sensor chips as easily as theZ_EGFR 1907’_ (Supplementary Figure S2A). The tight binding of Cetuximab to the EGFR sensor reduces the ability of the chip to reach equilibrium binding, so the K_D_ value determined in this study is not accurate. Our study determined Cetuximab to have an average K_D_ value of 5.3 pM ± 9.9 SD for sEGFR1-_621_ which is higher affinity than the reported K_D_ value of ∼1nM for Cetuximab binding to EGFR(57). There was a significant difference between K_on_ and K_off_ values calculated for both forms of the sEGFRs under local 1:1 binding model fits (Table S3). It is possible that upon binding of Cetuximab to the EGFR surface, the conformation of the EGFR changes from a low affinity receptor to a high affinity binding state (Table 1). X-ray crystallographic and binding studies have shown that Erbitux (Cetuximab) binds exclusively to domain III of soluble extracellular region of the EGFR (sEGFR)(51, 58).

A competition screen between Z_EGFR 1907’_ affibody and Cetuximab was measured using an A-B-A assay format where a flanking solution was injected before and after the sample. We used this assay to investigate the binding site of the Z_EGFR 1907’_ affibody to the EGFR. As shown in Figure 2B and Figure S1B binding of Cetuximab blocks binding of Z_EGFR 1907’_ affibody to sEGFR_1-621._ The slight dip in response units upon injection of the second component (B: Z_EGFR 1907’_ affibody) to Cetuximab suggests occurrence of some non-specific binding as Cetuximab is not expected to come off within 400 sec. Also, the relatively fast off rate for Z_EGFR 1907’_ affibody allows Cetuximab to bind to newly unbound sEGFR_1-621_ sites, however when Cetuximab is mixed with Z_EGFR 1907’_ affibody in excess, this does not occur, indicating direct competition between the EGFR binders. Together these data show that the Z_EGFR 1907’_ affibody shares the Cetuximab binding site (i.e. EGFR-domain III).

### EGFR expression and cell surface availability in colorectal and breast cancer cell lines

EGFR is overexpressed in majority of (50-80%) of CRCs(59) and these elevated levels of expression have been linked with tumor aggression and poor survival(14, 59). Cancer cells overexpressing EGFR appear to be ideal candidates for new targeted therapeutics such as the affibody polyplexes, but was not clear that all the EGFR would be on the cell surface and accessible to the polyplexes. Consequently, we compared the published proteomic data, i.e. total EGFR levels(42) with the cell surface EGFR levels measured by flow cytometry for a range of CRC cell lines and some breast cancer cell lines. Cell lines with increased EGFR expression from the proteomic analyses also had more cell surface EGFR (Figure 3A, Spearman correlation coefficient (r_s_) 0.74). For example, Difi cells have the highest levels of EGFR expressed on the cell surface (Figure 3A). A small number of CRC cells, e.g. SW620 and Colo201 and SW480 showed little to no EGFR expression in the proteomic or FACS studies. Thus, we used the SW620 cell line as an EGFR negative control in our experiments.

**Figure 3.**
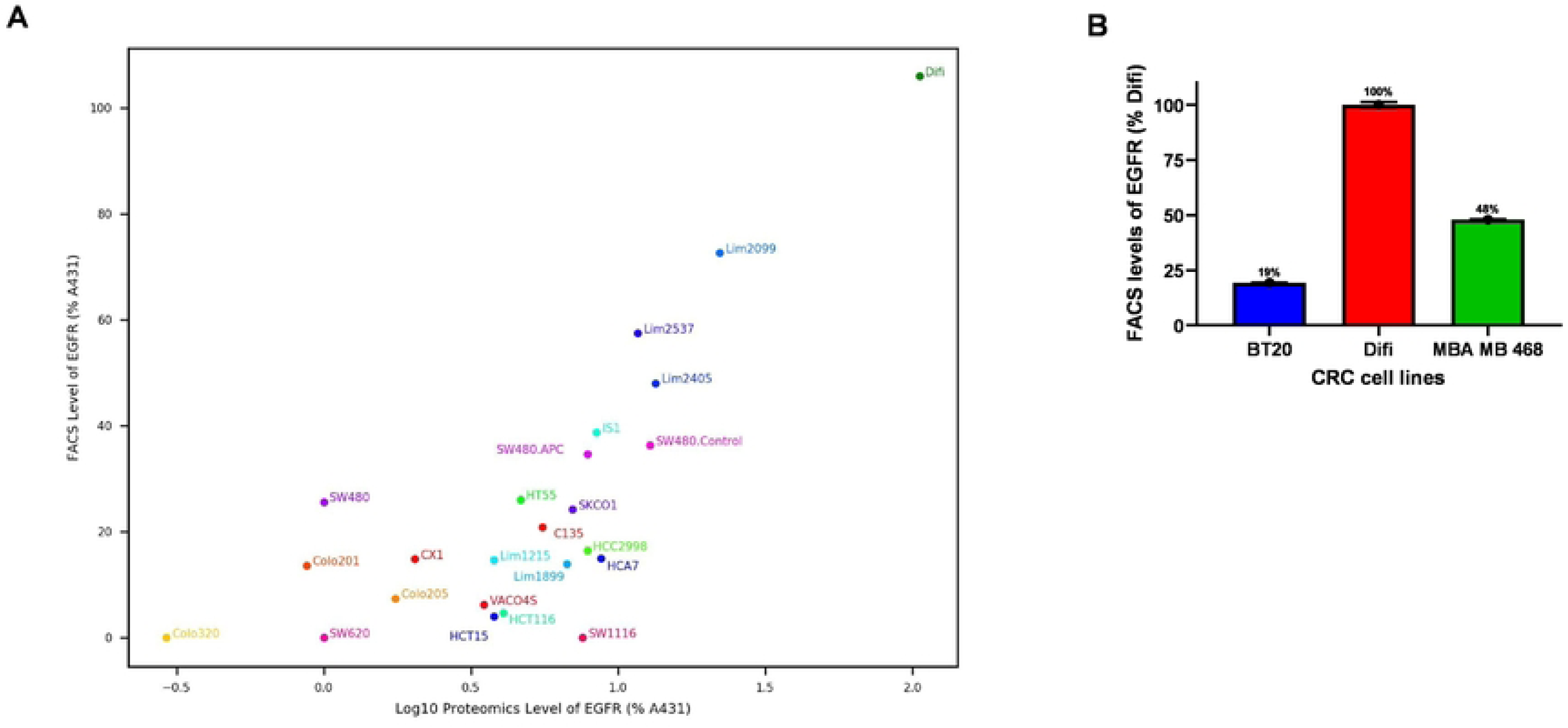
Relationship between cell surface EGFR and total EGFR in cancer cell lines. **(A)** Cell lines were incubated with anti-EGFR-PE and the PE fluorescence was measured by flow cytometry. The median fluorescence intensity for the respective quadrant was used to calculate relative levels of EGFR expression on cell surface compared to the A431 cell line. Proteomic data was calculated from mass spectrometer peptide counts. **(B)** Relative levels of EGFR expression on cell surface of breast cancer cell lines, BT-20 and MDA-MB-468, compared to the Difi cell line.

Two triple-negative breast cancer (TNBC) cell lines, BT-20 and MDA-MB-468 overexpress EGFR(57, 60). At present, TNBC patients have limited options for treatment and we expected that the EGFR-directed polyplexes might be suitable for treating TNBC patients whose tumors overexpress the EGFR. Cell surface expression of EGFR levels on these cell lines was determined. Both breast cell lines have lower levels of surface EGFR compared to the Difi cell line. MDA-MB-468 has slightly less than half the Difi EGFR levels (48%) and the BT-20 cell line had approximately a quarter of the Difi EGFR levels (19%) (Figure 3B). Similar results have been reported for total EGFR levels using Western blot analysis (Mohan *et al.*, 2021). The BT-20 and MDA-MB-468 EGFR levels are similar to the EGFR levels on CRC cell lines C135 and Lim2405 respectively (see Figure 3A,3B). Thus, both BT-20 and MDA-MB-468 cell lines can be considered to have moderate levels of EGFR on the cell surface.

### *In vitro* cancer cell killing by PPEA-polyplexes

To assess the cytotoxic activity of the PPEA-polyplex on cells in culture, we used cell lines that expressed different levels of the receptor (from undetectable to more than 10^6^ EGFR per cell). After preparation of the PPEA-polyplex and before use, we checked the size of the polyplex using the zetasizer. The particle size of the PPEA-polyplex was 119 ± 26nm. The PPEA triconjugate (i.e. the affibody-PEI-PEG complex without the polyIC) was used as a control for nonspecific toxicity to the cells. As illustrated in Figure 4A and 4B, the PPEA-polyIC-polyplex induced a strong killing of EGFR-over-expressing cells (1-2x10^6^ EGFRs/cell) as compared to a cell line devoid of EGFR (U138-MG). In contrast, treatment with the PPEA, polyI-polyplex did not result in cell killing in either of the cell lines tested (Figure 4C), indicating that growth inhibitory activity induced by PPEA transfection is polyIC-dependent. PPEA-polyIC-polyplexes also produced a significant growth inhibitory effect in the SK-BR-3, BT-474 and MDA-MB-231 cell lines, which express medium levels of the receptor (1x10^5^-3x10^5^ EGFRs/cell) (Figure 5). These results show that PPEA-polyIC-polyplexes inhibit the proliferation of tumor cell lines with medium to high levels of EGFR expression.

**Figure 4.**
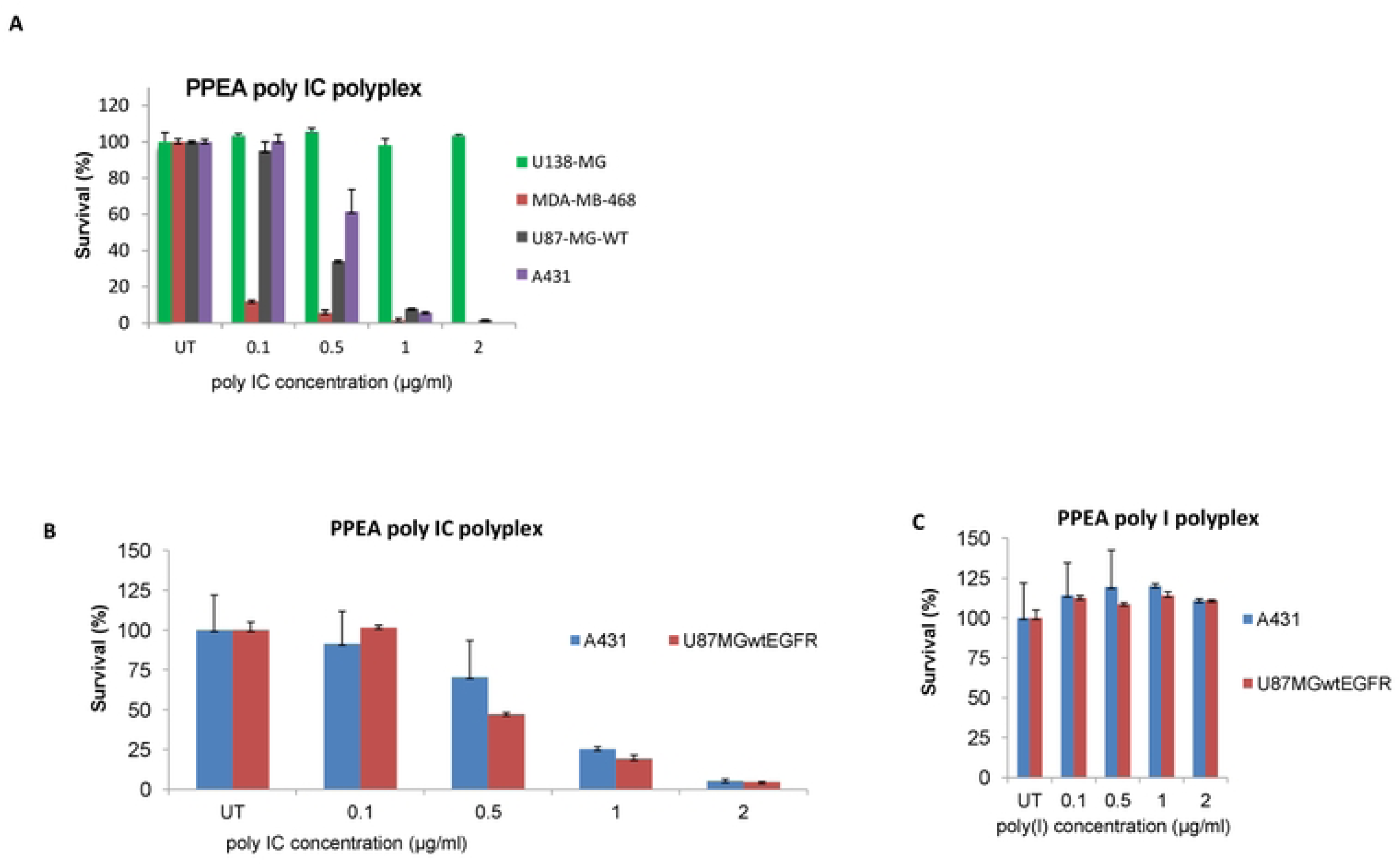
Anti-tumor activity of the PPEA-polyIC-polyplex *in vitro*. **(A)** PPEA-polyplexes selectively kill cell lines overexpressing EGFR. Cells were seeded in duplicates into 96-well plates at a density of 5000 cells in 0.1 ml medium per well and grown overnight. Cells were then transfected with polyIC at the indicated concentrations using the PPEA complex. PEI-PEG ratio=1:1; w/w ratio PEI: polyIC =0.78. U138MG cells do not express EGFR; U87MGwtEGFR cells express 1x10^6^, A431 express 2-3X10^6^ and MDA-MB-468 express 2x10^6^ EGFRs/cell. **(B,C**) *In vitro* anti-tumor activity of PPEA-polyIC-polyplex is polyIC-specific. A431 and U87MGwtEGFR cell lines were transfected with same doses of PPEA-polyIC-polyplex (**B)** or PPEA polyI polyplex (**C**), which served as negative control. Viability was measured by the PrestoBlue Cell Viability Reagent (Invitrogen), according to the manufacturer’s instructions, at 72 hours after transfection. These experiments were repeated three times with a representative experiment shown.

**Figure 5.**
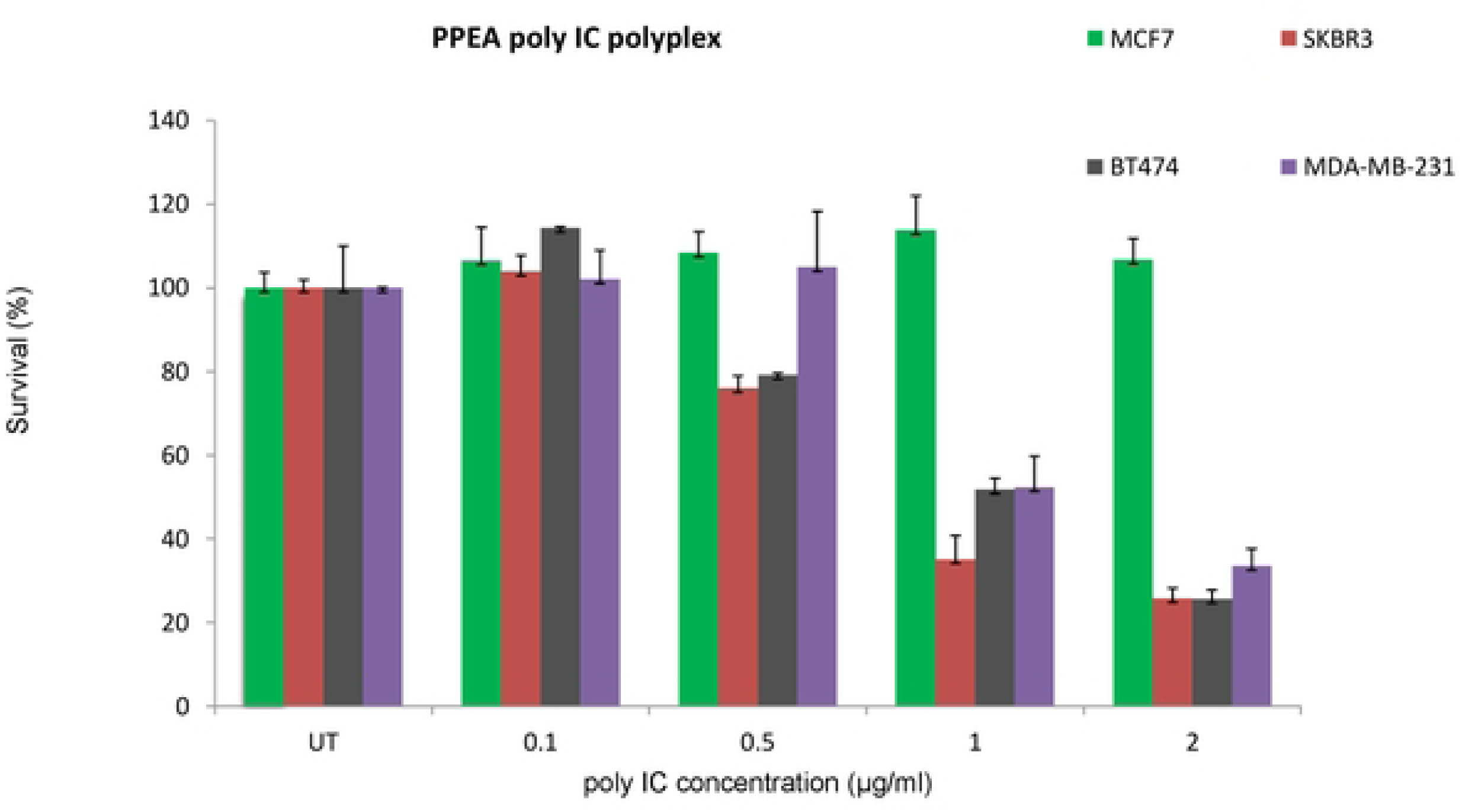
*In vitro* anti-tumor cell activity of the PPEA-polyplex in cell lines with medium levels of EGFR. Cells were transfected with polyIC at the indicated concentrations using PPEA. Cell viability was measured as described in Figure4. MDA-MB-231 cells express 2.5-3X10^5^ EGFRs/cell, SK-BR-3 express 3X10^5^, BT-474 express 10^5^ EGFRs/cell and MCF7 cells express 5x10^3^ EGFRs/cell.

To investigate the relationship between the level of EGFR expression on cell surface and the potency of the PPEA-polyplex, we determined EC_50_ values for two colon cancer cell lines SW480 and DiFi and two breast cancer cell lines BT-20 and MDA-MB-468. Upon treatment with PPEA-polyplex (Table 2, Figure S3), the CellTiter Glo assays showed significant growth inhibition of the CRC cell lines with medium levels of EGFR overexpression such as LIM1215 (EC_50_ < 300 ng/ml). Despite the very high levels of EGFR on DiFi cells the PPEA-polyplex was no more potent, than it was on the LIM1215 cells (Table 2, Figures S3B and S3C).

**Table 2.**
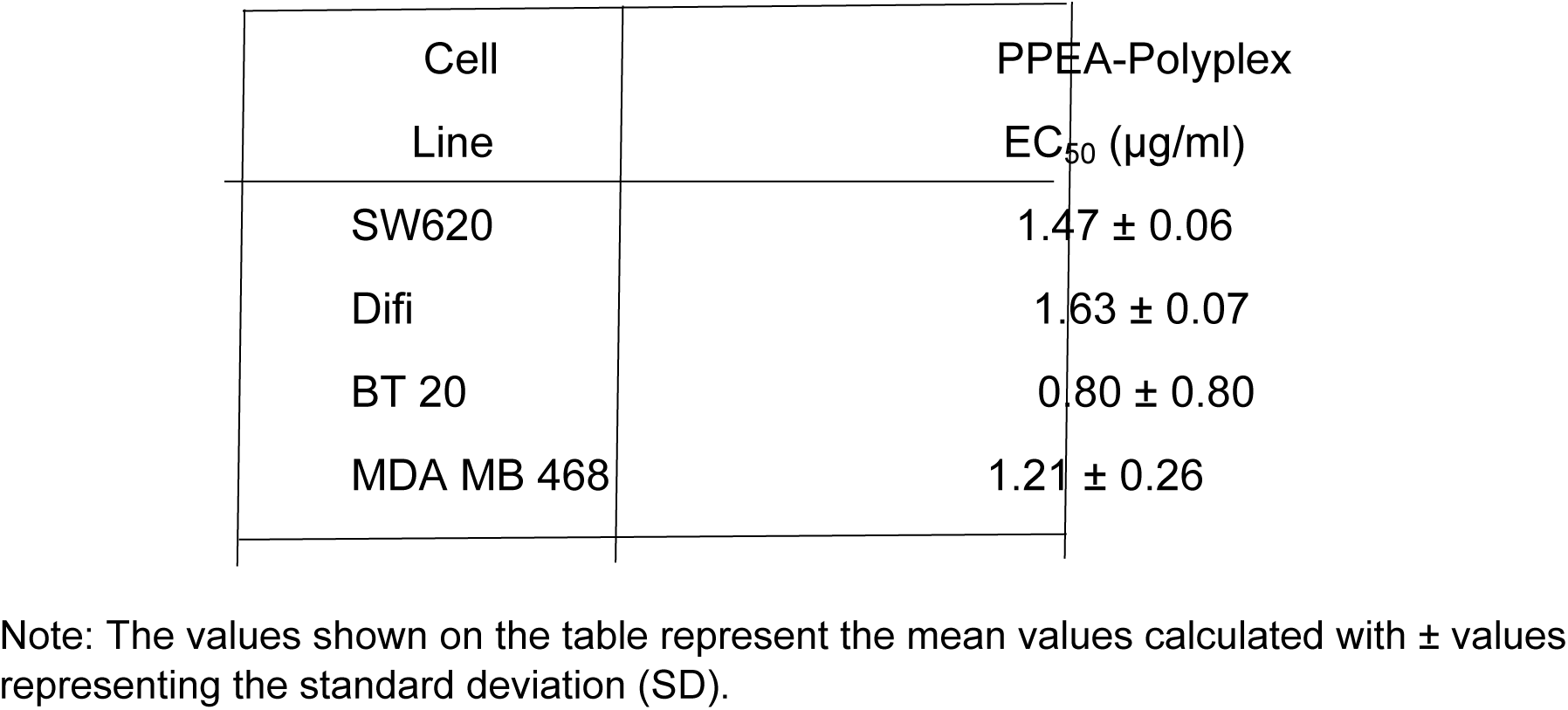
Cytotoxicity of the PPEA-polyplex on selected cancer cell lines.

| Cell<br>Line | PPEA-Polyplex<br>EC<sub>50</sub> (µg/ml) |
| --- | --- |
| SW620 | 1.47 ± 0.06 |
| Difi | 1.63 ± 0.07 |
| BT 20 | 0.80 ± 0.80 |
| MDA MB 468 | 1.21 ± 0.26 |

Note: The values shown on the table represent the mean values calculated with ± values representing the standard deviation (SD).

The PPEA-polyplex was a potent inhibitor of the breast cancer cell lines BT-20 and MDA-MB-468 even though they have less than half the EGFR levels of A431 cells, indeed the BT-20 cell line was 2 orders of magnitude more sensitive to PPEA-polyplex killing than the other cells (Table 2).

### PPEA-polyplex does not induce EGFR phosphorylation

We next examined whether the affibody retains its inability to phosphorylate EGFR, when tethered to PEI-PEG. We treated MDA-MB-468 cell with the PPEA-polyplex and analyzed the autophosphorylation of EGFR (Figure 6) and no EGFR autophosphorylation was detected in cells challenged with PPEA-polyplexes.

**Figure 6.**
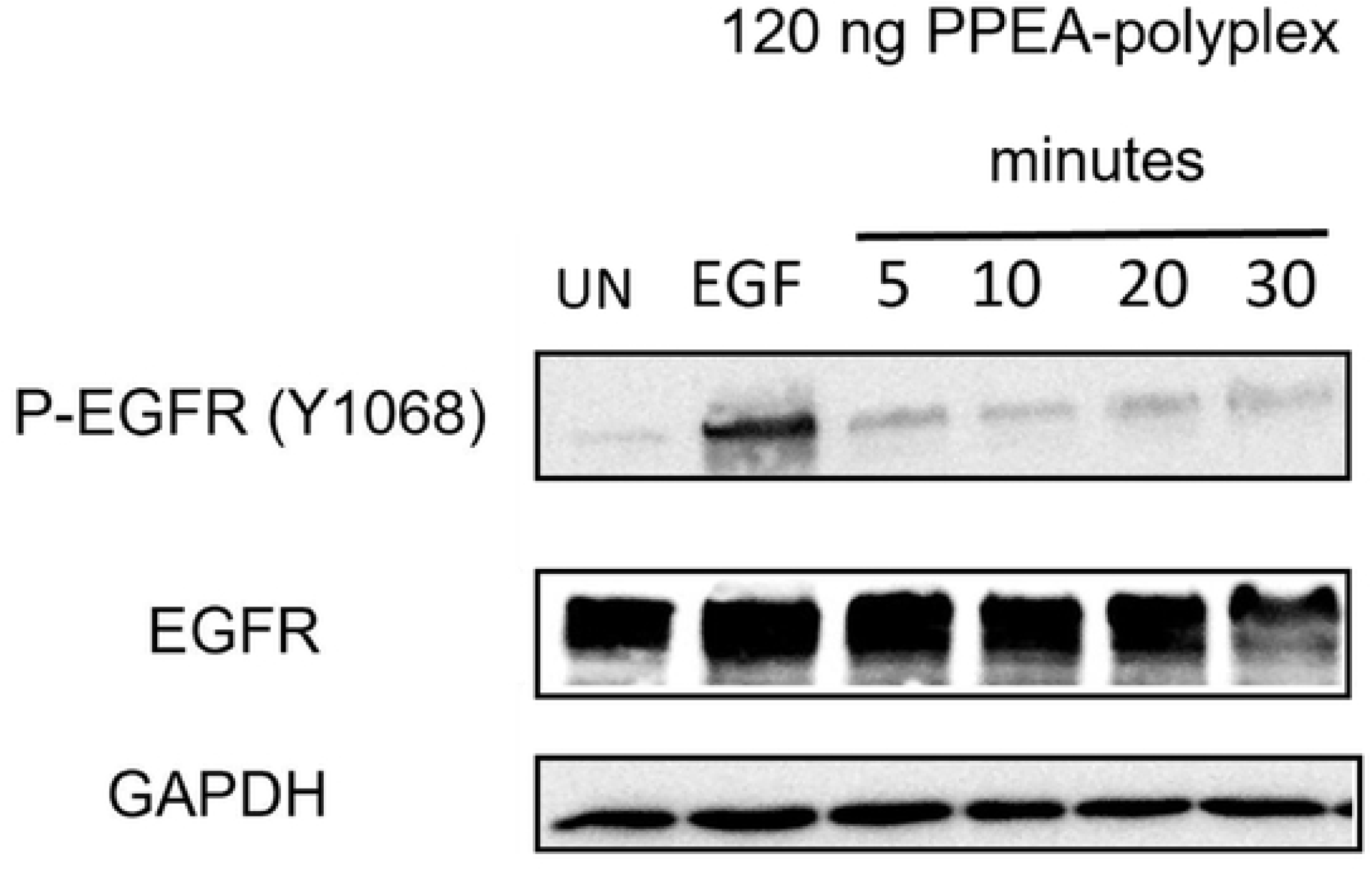
Western blot analysis of autophosphorylated and total EGFR in MDA-MB-468 cells upon addition of the PPEA-polyplex. Cells were treated as described in “Material and methods”. Untreated cells (UN) were used as negative control and cells incubated with EGF for 5 minutes were used as the positive control.

### The PPEA-polyplex induces expression of chemotactic cytokines in cancer cells expressing high and moderate levels of EGFR

Tumor cells that internalize polyIC are induced to secrete cytokines that attract immune cells(61, 62). In our previous studies, we showed that polyIC targeted by PPE, induced expression of Interferon ɣ-induced protein 10 kDa (IP-10), and growth-regulated protein α (Gro-α) in EGFR-overexpressing cell lines, but not in cells devoid of EGFR(23) (24). Gro-α and IP-10 are chemokines that play an active role in the recruitment of leukocytes to the domain in which they are secreted(63, 64). To evaluate the ability of PPEA-polyplex to induce EGFR-expressing cells to secrete inflammatory cytokines, we collected cell culture supernatants 48 hours after exposure to the PPEA-polyplex and assessed for the presence of cytokines Gro-α and IP-10. As can be seen in Table 3, cells harboring high or moderate levels of EGFR secrete chemotactic cytokines following challenge with PPEA-polyplex. In contrast, no cytokine secretion was detected in U138-MG and MCF-7 cells that are devoid of (or express low levels of) the EGFR, respectively, following treatment with the PPEA-polyplex. Interestingly, we observed significantly higher amounts of IP-10 in supernatants collected from MDA-MB-468, SK-BR-3 and BT-474 cells, compared to A431 and MDA-MB-231 (Table 3). This observation points to possible differences in signalling mechanisms mediated by PPEA-polyplex treatment in the cell lines tested. Collectively, these data demonstrate that polyIC transfection using PPEA stimulates release of pro-inflammatory cytokines by tumor cells expressing high and moderate levels of EGFR.

**Table 3.** Secretion of chemokines after transfection with PPEA-polyplex.

|  | **Cytokines (pg/ml)\*\*** |  |  |  |
| --- | --- | --- | --- | --- |
| | IP-10 | | Gro-α | |
|  | - | + | - | + |
| **PPEA-polyplex** |  |  |  |  |
| U138MG | 0 | 0 | 0 | 0 |
| MCF7 | 0 | 0 | 0 | 0 |
| A431 | 30±4 | 91±3 | 0 | 111±20 |
| MDA-MB-231 | 15±7 | 93.5±2 | 42±1 | 147±5 |
| SKBR3 | 73±13 | 634±35 | 0 | 142±21 |
| BT474 | 65±2 | 574±46 | 18±5 | 49±2 |
| MDA-MB-468 | 0 | 377±26 | 100±4 | 0 |

\*\*Cytokine levels measured by ELISA following 48 hours transfection using PPEA formulated with 2 μg/ml polyIC.

### PPEA-polyplex treated cancer cells activate human immune cells *in vitro*

We used peripheral blood mononuclear cells (PBMCs) to assess the ability of cytokine-enriched medium derived from PPEA-polyplex treated tumor cells to stimulate the immune cells(27). PBMCs consist of several types of immune cells, including monocytes, macrophages, NK (natural killer) cells and T-cells. When stimulated, PBMCs produce an array of cytokines which can be conveniently quantified by ELISA(65).

We tested whether the chemotactic stimuli secreted by cells that internalized polyIC stimulated PBMC to migrate toward the secreting cells(66). Using chemotactic chambers (see Materials and Methods) we showed that the cytokine-enriched medium from A431 cells transfected with PPEA-polyplex stimulated chemotaxis of PBMCs (Figure 7). In contrast, medium of U138MG treated with PPEA-polyplex did not induce chemotaxis. Thus, PBMCs are attracted by medium from EGFR overexpressing cells, which have been treated with PPEA-polyplex.

**Figure 7.**
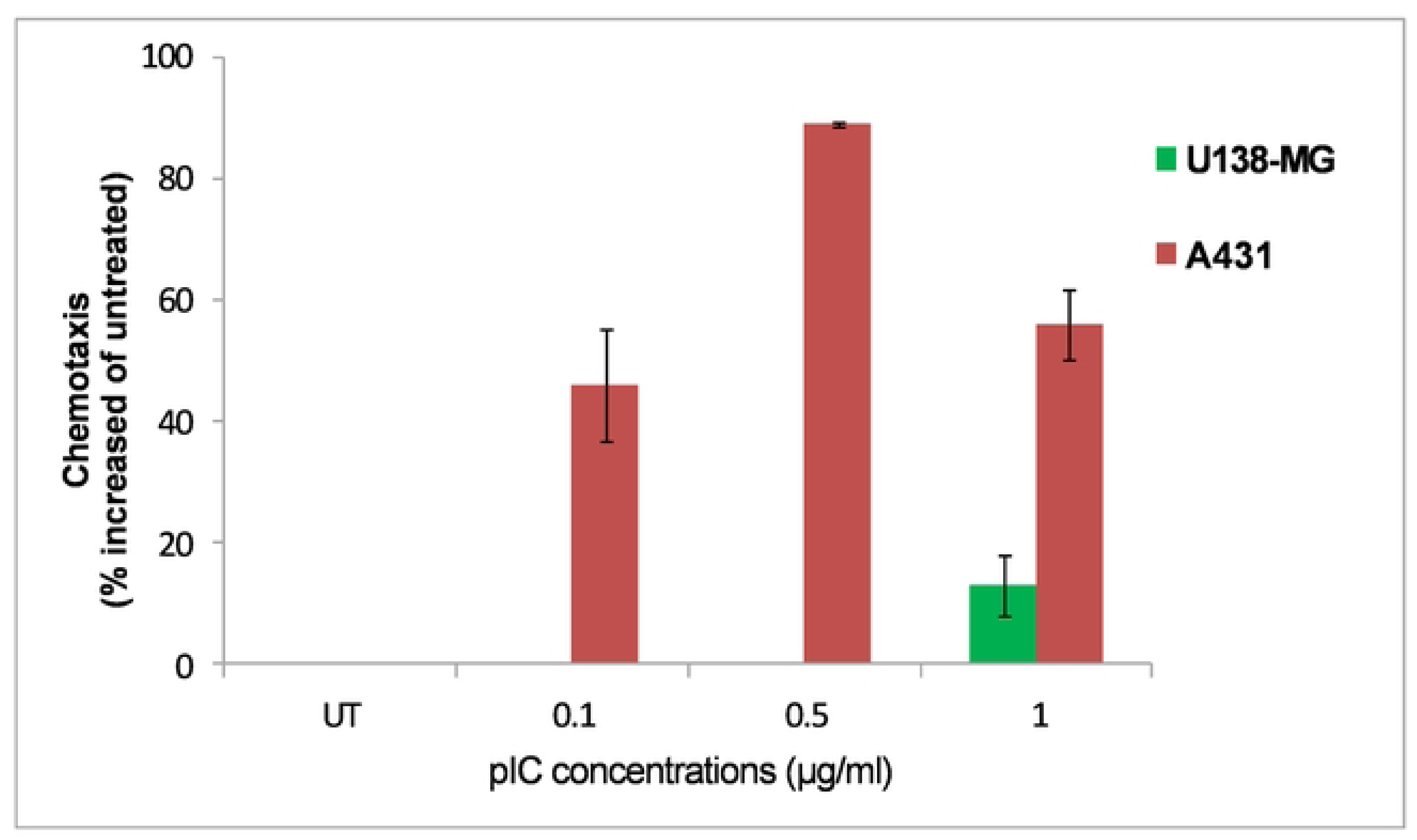
Comparison of anti-tumor activity by PPEA-polyIC-polyplex on PBMC chemotaxis *in vitro*. The PPEA-polyIC-polyplex selectively kills cell lines overexpressing EGFR. Cells were seeded in duplicates into 96-well plate at a density of 5000 cells in 0.1 ml medium per well and grown overnight. Cells were then transfected with polyIC at the indicated concentrations using the PPEA complexes. PEI-PEG ratio=1:1; w/w ratio PEI: Poly(IC) =0.78. U138MG cells do not express EGFR; U87MGwtEGFR cells express 1x10^6^, A431 express 2-3X106 and MDA-MB-468 express 2x10^6^ EGFRs/cell.

We examined the ability of the medium from PPEA-polyplex transfected cancer cells to stimulate PBMCs. Upon activation, PBMC produce cytotoxic cytokines, such as INF-ɣ and TNF-α, known to contribute to tumor suppression and eradication(67, 68). PBMC were challenged with culture supernatant from PPEA-polyplex transfected cells and their response was measured using ELISA. Table 4 illustrates the levels of INF-ɣ and TNF-α expression by the immune cells, detected 24 hours following the challenge. The data clearly show that the medium originated from polyIC-targeted A431, MDA-MB-231 and SK-BR-3 cells (polyIC, 2 µg/ml) stimulate the production of cytotoxic cytokines as compared to culture supernatant from untreated cells or unchallenged PBMC cells (Table 4). We conclude that the culture supernatants derived from polyIC-targeted EGFR overexpressing cells sustain the capability to selectively stimulate the immune cells whereas cells devoid of EGFR do not.

**Table 4.** Evaluation of cytotoxic cytokine levels secreted by activated PBMC cells.

| Cytokine (pg/ml)<sup>***</sup> |  |  |  |  |
| --- | --- | --- | --- | --- |
| PPEA polyplex | TNF-α | | INF-γ | |
|  | - | + | - | + |
| A431 | 98±24 | 560±26 | 330±15 | 73±7 |
| MDA-MB-231 | 71±10 | 311±58 | 7±1 | 126±21 |
| SKBR3 | 57±7 | 147±9 | 10±6 | 52±14 |
| PBMC<sup>a</sup> | 44±6 | | 9±2 | |

MDA-MB-231, A431 and SKBR-3 cell lines were seeded in duplicates into 96-well plate at a density of 10,000 cells in 0.1 ml per well. Cells were then transfected with the PPEA-polyplex (PPEA formulated with 0.5-2 μg/ml of polyIC). Following 48 hours of incubation, 200 μl of supernatants from each cell line were collected and added to PBMCs seeded in duplicates into 96-well plate at a density of $3 \times 10^6$ in 0.1 ml medium per well. Following 24 hours incubation, the supernatants were collected and subjected to TNF-α and INF- $\gamma$ determination using ELISA (PeproTech). Data represent means of duplicate wells ± SD and are representative of 2-3 experiments.
<sup>\*\*\*</sup>Cytokine secretion measured by ELISA following exposure of PBMCs to medium from cells transfected using PPEA formulated with 2 μg/ml polyIC.
<sup>a</sup> Shows expression of cytokines in PBMCs incubated with growth medium alone and served as a negative control.

### PBMC-mediated bystander effect

Data from clinical studies has demonstrated a correlation between the presence of tumor infiltrating lymphocytes which produce cytokines such as INF-ɣ and TNF-α, and better outcomes for patients with many different cancers(69, 70). Previously, we reported that expression of INF-ɣ and TNF-α by PBMCs strongly enhanced the bystander killing of untransfected tumor cells(27). To assess the PBMC-mediated bystander effect, MDA-MB-231 and SK-BR-3 cells were first transfected with PPEA-polyplex and 24 hours later, freshly isolated PBMCs were added to the transfected cells for co-incubation. Following a further 24 hours, medium from the transfected cells co-cultured with PBMC, was added to newly seeded, untreated cells (Figure 8A). The PBMC-mediated bystander killing was examined 72 hours later, and compared with the “direct” bystander killing, mediated by medium from PPEA-polyplex-treated MDA-MB-231 and SK-BR-3 cells (Figure 8B and 8C). The medium derived from polyIC-transfected cells co-incubated with PBMCs, exerted a strong PBMC-mediated bystander effect, killing up to 80-90% of the nontransfected cells (Figure 8C). Medium derived from MDA-MB-231 or SK-BR-3 cells challenged with PPEA-polyplex alone, eliminated only 20 and 35 % of untreated tumor cells, respectively (Figure 8B). These results demonstrate that PPEA-polyplex treatment and immune cells display a cooperative cell-killing activity by enhancing bystander killing of EGFR-overexpressing tumors.

**Figure. 8.**
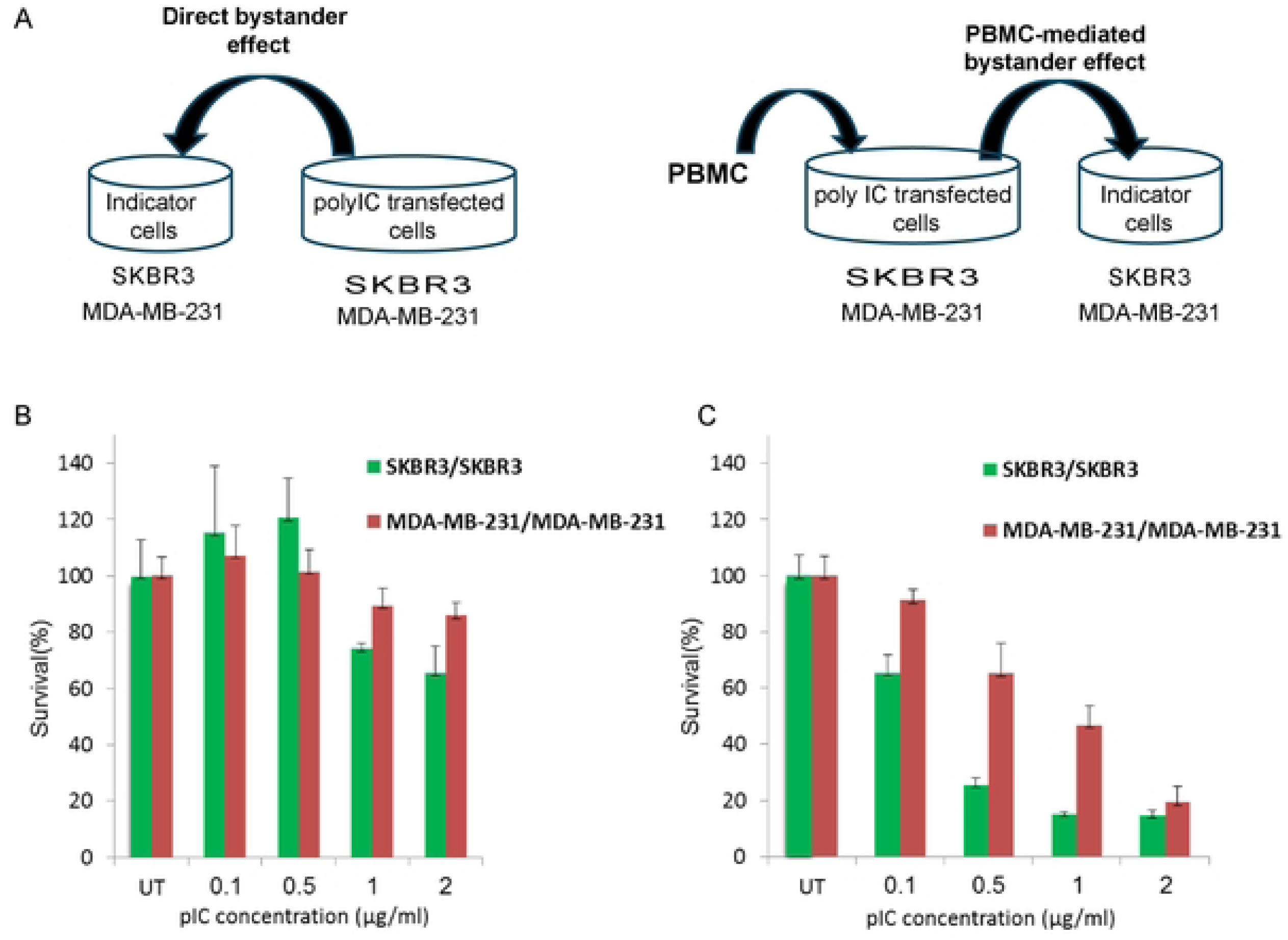
“Direct” versus PBMC-mediated bystander effect. 50,000 of MDA-MB-231 and SK-BR-3 cells were seeded into 24-well plates in duplicates and grown overnight with 1 ml medium per well. Cells were then transfected with the PPEA-polyplex at the indicated concentrations. To test the “direct” bystander effect, 48 hours following transfection 0.1 ml of medium from challenged cells was exchanged for 0.1 ml medium from non-transfected cells (“indicator cells) seeded on 96 well plates (4000 cell/well) 24 hours earlier. To analyze the PBMC-mediated bystander effect, 5x105 of PBMCs were added to transfected cells 24 hrs following transfection. After 24 hrs of co-culture, 0.15 ml of medium from the “conditioned medium” was added to untreated cells seeded into 96 well plates (4000 cell/well) 24 hours earlier. Survival of untransfected cells was determined by methylene blue assay, 72 hrs after challenge with the conditioned medium. (A) Shows experiment design. (B) Shows the “direct” bystander effect of medium from polyIC-targeted MDA-MB-231 and SK-BR-3 cells on unchallenged MDA-MB-231 and SK-BR-3, respectively. (C) Shows PBMC-mediated bystander effect of PBMCs co-incubated with polyIC-transfected MDA-MB-231 and SK-BR-3 cells on unchallenged MDA-MB-231 and SK-BR-3, respectively.

### PPEA complexes induce bystander effects

“Direct” versus PBMC-mediated bystander effects were also measured by transfecting MDA-MB-231 and SK-BR-3 cells with the PPEA-polyplex and testing the direct or indirect bystander effects of the medium on PBMCs. Supplementary Figure S4A shows the experimental design, Supplementary Figure S4B shows the “direct” bystander effect of medium from polyIC-targeted MDA-MB-231 and SK-BR-3 cells on unchallenged MDA-MB-231 and SK-BR-3, respectively and Figure S4C the PBMC-mediated bystander effect of PBMCs co-incubated with polyIC-transfected MDA-MB-231 and SK-BR-3 cells on unchallenged MDA-MB-231 and SK-BR-3, respectively. The supernatants from PPEA-polyplex treated cells activate human immune cells.

### Efficacy of PPEA-polyplexes on human A431 xenografts growing in nude mice

The *in vivo* antitumor activity of the PPEA-polyplex was examined using the A431 subcutaneous xenograft model. Figure 9 shows that the PPEA-polyIC-polyplex inhibited tumor growth strongly. The tumors treated with the PPEA-poly Inosine (polyI)-polyplex grew at a similar rate as tumors in the untreated group.

**Figure 9.**
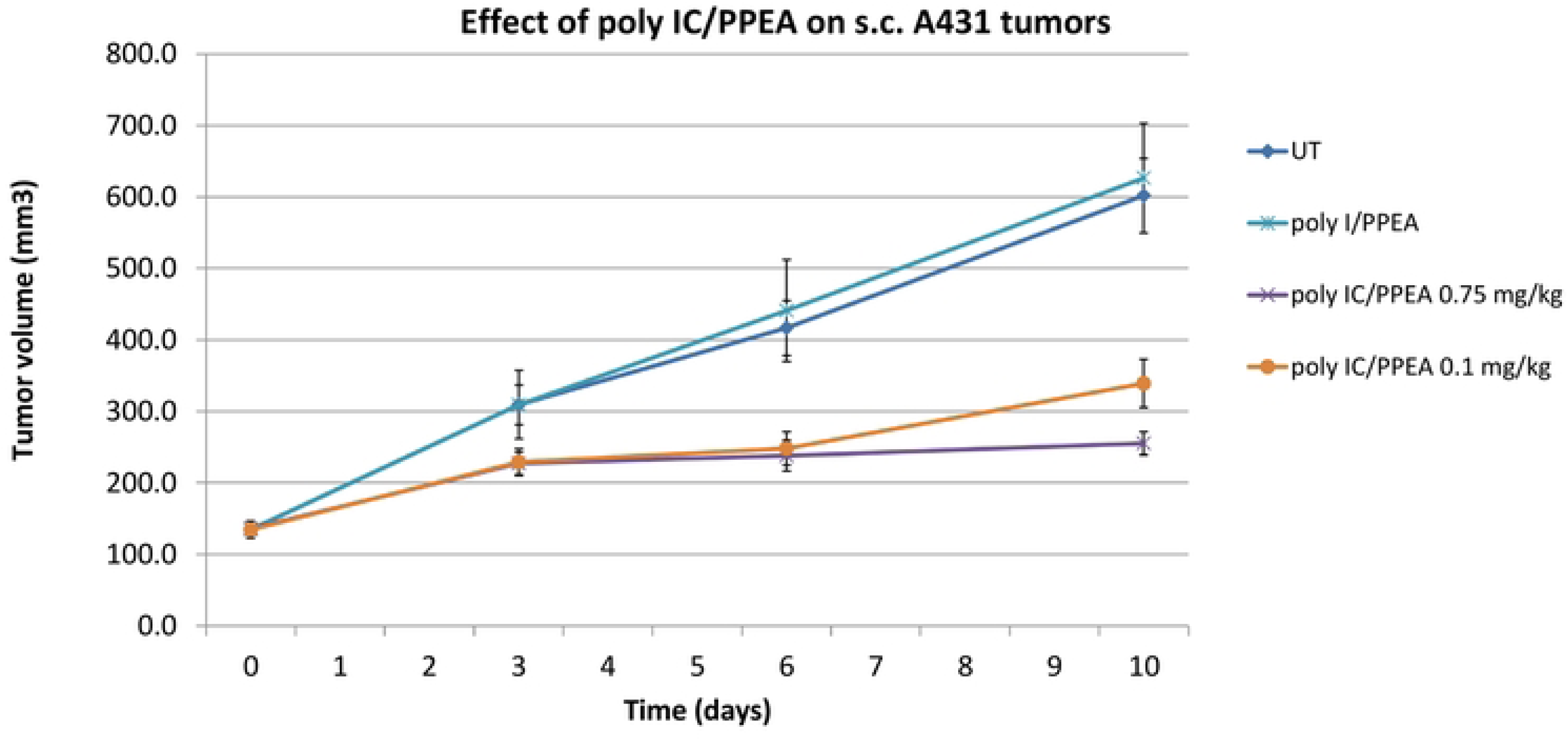
*In vivo* efficacy of PPEA-polyIC and poly I-polyplexes. Female nude mice were injected subcutaneously (s.c.) with A431 cells as described in the Methods section. After tumor establishment, mice were randomly divided into five groups and treated with daily intravenous injections (6 days/week) of the indicated polyplex at either 0.75 mg/kg or 0.1 mg/kg for 10 days, totaling 9 injections. Tumor volumes were measured on days 3, 6 and 10, and the chart presents the average tumor volume for each group, with error bars representing the standard error of the mean (SEM).

## Discussion

Combining the immune system’s ability to kill cancer cells together with engineered antibodies that target cancer cells has been shown to be an effective treatment of some cancers(71–75). Signaling from the EGFR family has been reported to drive the growth of many human cancers(13, 76–80). This motivated us to target the EGFR and at the same time attempt to stimulate anti-tumor immune responses and activate killing pathways within the tumor cells. Targeted polyIC therapy offers a promising approach for improving cancer therapy (78–80). Chemical vectors for therapeutic applications have the advantage over viral vectors since they can be custom designed to target specific cells, are less immunogenic and cannot interact with the host genome. Affibodies directed against the EGFR or EGFR ligands coupled to a scaffold which can bind polyIC have the potential to deliver polyIC to cancers with elevated, active EGFR.

We produced and purified the Z_EGFR 1907’_-affibody-polyIC-polyplex (PPEA-polyplex). The efficiency of drug delivery is influenced by the biophysical properties of the polyplex such as size, shape and surface charge(81, 82). Previous biophysical and structural studies have revealed that sEGFR_1-501_ (domain I, II and III) binds to EGF ligand with greater affinity than the full-length EGFR ectodomain(83). Cetuximab (monoclonal antibody) interacts exclusively with domain III sEGFR, hindering ligand binding and sterically preventing dimerization(51). Analysis of the EGFR binding of the PPEA-polyplex revealed an average K_D_ of 6.74 nM to sEGFR_1-621,_ suggesting domain IV may function to stabilize binding of the affibody (Figure 2). Kinetic data models together with the significant difference in the k_on_ and k_off_ values suggest the presence of two binding site conformations for theZ_EGFR 1907’_ on sEGFR_1-621_, with one binding site much weaker than the other. This may be due to the presence of both tethered/non tethered forms of sEGFR_1-621_ or the orientation of the sEGFR_1-621_ on the sensor chip. Higher K_D_ values obtained for sEGFR_1-501_ is most likely to be due to its inability to form the untethered back-to-back conformation which occurs on dimerization and which is required for tight binding of Z_EGFR 1907’_-affibody. The high affinity binding of Cetuximab to sEGFR makes it difficult to measure the K_D_ values using the biosensor. However, the trend in K_D_ values suggests there was much tighter binding between Cetuximab and sEGFR_1-621_ than between Cetuximab and sEGFR_1-501_, indicating the need for domain IV to facilitate tight binding of Cetuximab to the sEGFR_1-621._

Saturated binding of Cetuximab followed byZ_EGFR 1907’_ shows that Z_EGFR 1907’_ has minimal available space to bind, whilst the opposite is true when the affibody dissociates and thus providing binding sites for Cetuximab. These findings lead us to believe that the affibody binds to the same location on the EGFR as Cetuximab (Erbitux) – i.e. domain III.

The potency of these polyplexes was tested on a range of selected cell lines expressing different levels of EGFR: CRC cell lines, breast cancer cell lines and A431 cell line (Figure 4). In general cells with more receptors are killed more effectively by the EGF-conjugated polyplex, however, some cells with high levels of EGFR are still resistant to the polyplex, whereas other cells with moderate levels of EGFR are sensitive to low concentrations of the polyplex. For example, the breast cancer cell lines used in this study have been reported to have only moderate levels of EGFR(57, 60) yet these cell lines were shown to be the most sensitive to the polyplexes. There are several possible explanations, e.g. EGFR receptor turnover might be required, or cell survival pathways are deficient in some cells. Further investigation is required to identify biomarkers which can indicate which patient tumors are likely to respond to the polyplex. It will be important that each patient’s tumor is tested for EGFR levels and that organoids from their tumor can be assayed to determine their likely responses to the polyplexes. It will be interesting to investigate whether the rate of turnover of EGFR in these cell lines plays a part in their susceptibility to the polyplex.

We tested the potency of polyIC complexed with a non-viral vector PPEA - an EGFR-homing vector which does not activate the EGFR (Figure 5). Previous studies had shown that GE11, an EGFR-homing peptide devoid of receptor activation, showed good potency in both in cells and *in vivo*(84). Yet the affinity of the GE11 based vector is not sufficiently high to enable its development as a clinical candidate [32]. In contrast, the EGFR-binding affibody molecule (Z_EGFR1907’_) exhibits nanomolar binding affinity to EGFR, does not activate the kinase moiety of the receptor(85, 86) but its polyIC-polyplex (PPEA) kills EGFR positive tumor cells effectively.

Significantly, we found that PPEA-polyplexes induced a strong EGFR-specific killing effect in breast cell lines expressing medium levels of the receptor (1x10^5^-3x10^5^ EGFRs/cell (Figure 5). It is possible that the high efficacy of polyIC delivery by PPEA, along with its inability to activate EGFR, is the origin of the efficacy of PPEA-polyplexes in cells with moderate expression of the receptor. The ability of polyIC targeted by PPEA to induce expression of chemotactic cytokines in EGFR-overexpressing cells was comparable to that exhibited by an equivalent polyplex vector which used EGF target and activated the EGFR. Moreover, the supernatant fluids from PPEA-polyplex treated cells were able to activate human immune cells and to induce production of cytotoxic cytokines.

*In vivo* antitumor activity of PPEA was determined using A431 subcutaneous xenografts (Figure 9). Based on the data gathered, it can be said that there was greater than a 7 days delay in the growth of the tumors in mice treated with PPEA-polyplexes. Human BT-20 cells would be expected to be killed in xenografts, but the immune deficient mice do not provide the immune responses which are likely to occur in patients.

In conclusion, PPEA polyplexes effectively eliminate tumors with moderate EGFR expression, though tumors with very high EGFR expression may remain resistant. They induce immune cytokines and promote bystander killing of tumors *in vitro*, without activating EGFR kinase activity. These findings underscore the therapeutic potential of PPEA polyplexes and warrant further evaluation in preclinical and clinical models.

## Acknowledgements

The authors wish to acknowledge Shoshana Klein for encouragement throughout this project and specifically for helping with the drafting of this manuscript. We are most grateful to Christoph Grohmann who provided expert guidance as we prepared the conjugates. Our early discussions with Donna Jovin and Thomas Jovin helped us formulate the project and they provided the initial affibody clones. Esteban Pombo-Villar at TargImmune Corporation also provided help with the development of this manuscript.

## Figure Legends

For Supplementary Tables (S1 -S3), Supplementary Figures (S1-S4) and Supplementary Figure Legends (S1-S4)

see separate file

## References

1. Carpenter G, Stoscheck CM, Preston YA, DeLarco JE. Antibodies to the epidermal growth factor receptor block the biological activities of sarcoma growth factor. Proceedings of the National Academy of Sciences. 1983;80(18):5627–30.

2. Liberman T, Nusbaum H, Razon N, Kris R, Lax I, Soreq H, et al. Amplification, enhanced expression of possible rearrangement of EGF receptor gene in primary human brain tumor of glial origin. Nature. 1985;313:144–7.

3. Carlsson J, Wester K, De La Torre M, Malmström P-U, Gårdmark T. EGFR-expression in primary urinary bladder cancer and corresponding metastases and the relation to HER2-expression. On the possibility to target these receptors with radionuclides. Radiology and oncology. 2015;49(1):50.

4. Saeki T, Salomon DS, Johnson GR, Gullick WJ, Mandai K, Yamagam K, et al. Association of epidermal growth factor-related peptides and type I receptor tyrosine kinase receptors with prognosis of human colorectal carcinomas. Japanese journal of clinical oncology. 1995;25(6):240–9.

5. Salomon DS, Brandt R, Ciardiello F, Normanno N. Epidermal growth factor-related peptides and their receptors in human malignancies. Critical reviews in oncology/hematology. 1995;19(3):183–232.

6. Kalyankrishna S, Grandis JR. Epidermal growth factor receptor biology in head and neck cancer. Journal of clinical oncology. 2006;24(17):2666–72.

7. Okabe T, Okamoto I, Tamura K, Terashima M, Yoshida T, Satoh T, et al. Differential constitutive activation of the epidermal growth factor receptor in non–small cell lung cancer cells bearing EGFR gene mutation and amplification. Cancer research. 2007;67(5):2046–53.

8. Hashmi AA, Naz S, Hashmi SK, Irfan M, Hussain ZF, Khan EY, et al. Epidermal growth factor receptor (EGFR) overexpression in triple-negative breast cancer: association with clinicopathologic features and prognostic parameters. Surgical and Experimental Pathology. 2019;2(1):1–7.

9. Pouliot N, Nice EC, Burgess AW. Laminin-10 mediates basal and EGF-stimulated motility of human colon carcinoma cells via α3β1 and α6β4 integrins. Experimental cell research. 2001;266(1):1–10.

10. Sizeland AM, Burgess AW. Anti-sense transforming growth factor alpha oligonucleotides inhibit autocrine stimulated proliferation of a colon carcinoma cell line. Molecular biology of the cell. 1992;3(11):1235–43.

11. Porta R, Sanchez-Torres J, Paz-Ares L, Massuti B, Reguart N, Mayo C, et al. Brain metastases from lung cancer responding to erlotinib: the importance of EGFR mutation. European Respiratory Journal. 2011;37(3):624–31.

12. Kato S, Okamura R, Mareboina M, Lee S, Goodman A, Patel SP, et al. Revisiting epidermal growth factor receptor (EGFR) amplification as a target for anti-EGFR therapy: analysis of cell-free circulating tumor DNA in patients with advanced malignancies. JCO precision oncology. 2019;3:1–14.

13. Saletti P, Molinari F, De Dosso S, Frattini M. EGFR signaling in colorectal cancer: a clinical perspective. Gastrointest Cancer. 2015;5:21–38.

14. Spano J-P, Lagorce C, Atlan D, Milano G, Domont J, Benamouzig R, et al. Impact of EGFR expression on colorectal cancer patient prognosis and survival. Annals of oncology. 2005;16(1):102–8.

15. Xu MJ, Johnson DE, Grandis JR. EGFR-targeted therapies in the post-genomic era. Cancer and Metastasis Reviews. 2017;36(3):463–73.

16. Lievre A, Bachet J-B, Le Corre D, Boige V, Landi B, Emile J-F, et al. KRAS mutation status is predictive of response to cetuximab therapy in colorectal cancer. Cancer research. 2006;66(8):3992–5.

17. Gan HK, Walker F, Burgess AW, Rigopoulos A, Scott AM, Johns TG. The Epidermal Growth Factor Receptor (EGFR) Tyrosine Kinase Inhibitor AG1478 Increases the Formation of Inactive Untethered EGFR Dimers IMPLICATIONS FOR COMBINATION THERAPY WITH MONOCLONAL ANTIBODY 806. Journal of Biological Chemistry. 2007;282(5):2840– 50.

18. Blasco MT, Navas C, Martín-Serrano G, Graña-Castro O, Lechuga CG, Martín-Díaz L, et al. Complete regression of advanced pancreatic ductal adenocarcinomas upon combined inhibition of EGFR and C-RAF. Cancer cell. 2019;35(4):573–87. e6.

19. Saba NF, Chen ZG, Haigentz M, Bossi P, Rinaldo A, Rodrigo JP, et al. Targeting the EGFR and immune pathways in squamous cell carcinoma of the head and neck (SCCHN): forging a new alliance. Molecular cancer therapeutics. 2019;18(11):1909–15.

20. Goldstein M, Rudra S, Dahiya S, Tsien C, Huang J. Prognostic value of EGFR Amplification in Glioblastoma Patients treated with Radiation Therapy and Concurrent Temozolomide. International Journal of Radiation Oncology• Biology• Physics. 2019;105(1):E98–E9.

21. Sapoznik S, Hammer O, Ortenberg R, Besser MJ, Ben-Moshe T, Schachter J, et al. Novel anti-melanoma immunotherapies: disarming tumor escape mechanisms. Clinical & developmental immunology. 2012;2012:818214.

22. Ramos CA, Dotti G. Chimeric antigen receptor (CAR)-engineered lymphocytes for cancer therapy. Expert opinion on biological therapy. 2011;11(7):855–73.

23. Shir A, Ogris M, Wagner E, Levitzki A. EGF receptor-targeted synthetic double-stranded RNA eliminates glioblastoma, breast cancer, and adenocarcinoma tumors in mice. PLoS medicine. 2006;3(1):e6.

24. Broka D, Klein S, Shir A, Schade B, Saxena M, Dasargyri A, et al. Targeted apoptotic immune modulator for the treatment of metastatic EGFR-positive solid tumors. Proceedings of the National Academy of Sciences. 2025;122(22):e2500489122.

25. Kumar H, Koyama S, Ishii KJ, Kawai T, Akira S. Cutting edge: cooperation of IPS-1- and TRIF-dependent pathways in poly IC-enhanced antibody production and cytotoxic T cell responses. J Immunol. 2008;180(2):683–7.

26. Trumpfheller C, Longhi MP, Caskey M, Idoyaga J, Bozzacco L, Keler T, et al. Dendritic cell-targeted protein vaccines: a novel approach to induce T-cell immunity. Journal of internal medicine. 2012;271(2):183–92.

27. Shir A, Ogris M, Roedl W, Wagner E, Levitzki A. EGFR-homing dsRNA activates cancer-targeted immune response and eliminates disseminated EGFR-overexpressing tumors in mice. Clinical cancer research : an official journal of the American Association for Cancer Research. 2011;17(5):1033–43.

28. Benhar M, Engelberg D, Levitzki A. ROS, stress-activated kinases and stress signaling in cancer. EMBO reports. 2002;3(5):420–5.

29. Friedman M, Orlova A, Johansson E, Eriksson TL, Höidén-Guthenberg I, Tolmachev V, et al. Directed evolution to low nanomolar affinity of a tumor-targeting epidermal growth factor receptor-binding affibody molecule. Journal of molecular biology. 2008;376(5):1388– 402.

30. Feldwisch J, Tolmachev V, Lendel C, Herne N, Sjöberg A, Larsson B, et al. Design of an optimized scaffold for affibody molecules. Journal of molecular biology. 2010;398(2):232– 47.

31. Friedman M, Nordberg E, Hoiden-Guthenberg I, Brismar H, Adams GP, Nilsson FY, et al. Phage display selection of Affibody molecules with specific binding to the extracellular domain of the epidermal growth factor receptor. Protein engineering, design & selection : PEDS. 2007;20(4):189–99.

32. Gostring L, Chew MT, Orlova A, Hoiden-Guthenberg I, Wennborg A, Carlsson J, et al. Quantification of internalization of EGFR-binding Affibody molecules: Methodological aspects. International journal of oncology. 2010;36(4):757–63.

33. Brissault B, Kichler A, Guis C, Leborgne C, Danos O, Cheradame H. Synthesis of linear polyethylenimine derivatives for DNA transfection. Bioconjugate chemistry. 2003;14(3):581–7.

34. Brissault B, Kichler A, Leborgne C, Danos O, Cheradame H, Gau J, et al. Synthesis, characterization, and gene transfer application of poly (ethylene glycol-b-ethylenimine) with high molar mass polyamine block. Biomacromolecules. 2006;7(10):2863–70.

35. Joubran S, Zigler M, Pessah N, Klein S, Shir A, Edinger N, et al. Optimization of liganded polyethylenimine polyethylene glycol vector for nucleic acid delivery. Bioconjugate chemistry. 2014;25(9):1644–54.

36. Ungaro F, De Rosa G, Miro A, Quaglia F. Spectrophotometric determination of polyethylenimine in the presence of an oligonucleotide for the characterization of controlled release formulations. Journal of pharmaceutical and biomedical analysis. 2003;31(1):143–9.

37. Schaffert D, Kiss M, Rödl W, Shir A, Levitzki A, Ogris M, et al. Poly (I: C)-mediated tumor growth suppression in EGF-receptor overexpressing tumors using EGF-polyethylene glycol-linear polyethylenimine as carrier. Pharmaceutical research. 2011;28(4):731–41.

38. Goldenberg A, Masui H, Divgi C, Kamrath H, Pentlow K, Mendelsohn J. Imaging of human tumor xenografts with an indium-111-labeled anti-epidermal growth factor receptor monoclonal antibody. Journal of the National Cancer Institute. 1989;81(21):1616–25.

39. Trempe GL. Human breast cancer in culture. Recent results in cancer research Fortschritte der Krebsforschung Progres dans les recherches sur le cancer. 1976(57):33–41.

40. Huang HS, Nagane M, Klingbeil CK, Lin H, Nishikawa R, Ji XD, et al. The enhanced tumorigenic activity of a mutant epidermal growth factor receptor common in human cancers is mediated by threshold levels of constitutive tyrosine phosphorylation and unattenuated signaling. The Journal of biological chemistry. 1997;272(5):2927–35.

41. Lasfargues EY, Coutinho WG, Redfield ES. Isolation of two human tumor epithelial cell lines from solid breast carcinomas. Journal of the National Cancer Institute. 1978;61(4):967–78.

42. Wang J, Mouradov D, Wang X, Jorissen RN, Chambers MC, Zimmerman LJ, et al. Colorectal cancer cell line proteomes are representative of primary tumors and predict drug sensitivity. Gastroenterology. 2017;153(4):1082–95.

43. Grohmann C, Walker F, Devlin M, Luo M-X, Chüeh AC, Doherty J, et al. Preclinical small molecule WEHI-7326 overcomes drug resistance and elicits response in patient-derived xenograft models of human treatment-refractory tumors. Cell Death & Disease. 2021;12(3):268.

44. Berman HG. Sampling Distribution: Difference Between Means [Available from: https://www.stattrek.com/sampling/difference-in-means 2025, accessed 9/26/2025.

45. Reuveni H, Flashner-Abramson E, Steiner L, Makedonski K, Song R, Shir A, et al. Therapeutic destruction of insulin receptor substrates for cancer treatment. Cancer Res. 2013;73(14):4383–94.

46. Mizrachy-Schwartz S, Cohen N, Klein S, Kravchenko-Balasha N, Levitzki A. Up-regulation of AMP-activated protein kinase in cancer cell lines is mediated through c-Src activation. The Journal of biological chemistry. 2011;286(17):15268–77.

47. Elleman TC, Domagala T, McKern NM, Nerrie M, Lönnqvist B, Adams TE, et al. Identification of a determinant of epidermal growth factor receptor ligand-binding specificity using a truncated, high-affinity form of the ectodomain. Biochemistry. 2001;40(30):8930–9.

48. Bajaj M, Waterfield MD, Schlessinger J, Taylor WR, Blundell T. On the tertiary structure of the extracellular domains of the epidermal growth factor and insulin receptors. Biochimica et Biophysica Acta (BBA)-Protein Structure and Molecular Enzymology. 1987;916(2):220–6.

49. Lax I, Fischer R, Ng C, Segre J, Ullrich A, Givol D, et al. Noncontiguous regions in the extracellular domain of EGF receptor define ligand-binding specificity. Cell regulation. 1991;2(5):337–45.

50. Ward CW, Hoyne PA, Flegg RH. Insulin and epidermal growth factor receptors contain the cysteine repeat motif found in the tumor necrosis factor receptor. Proteins: Structure, Function, and Bioinformatics. 1995;22(2):141–53.

51. Li S, Schmitz KR, Jeffrey PD, Wiltzius JJ, Kussie P, Ferguson KM. Structural basis for inhibition of the epidermal growth factor receptor by cetuximab. Cancer cell. 2005;7(4):301–11.

52. Kozer N, Henderson C, Jackson JT, Nice EC, Burgess AW, Clayton AH. Evidence for extended YFP-EGFR dimers in the absence of ligand on the surface of living cells. Physical biology. 2011;8(6):066002.

53. Kozer N, Kelly MP, Orchard S, Burgess AW, Scott AM, Clayton AH. Differential and synergistic effects of epidermal growth factor receptor antibodies on unliganded ErbB dimers and oligomers. Biochemistry. 2011;50(18):3581–90.

54. Kozer N, Rothacker J, Burgess AW, Nice EC, Clayton AH. Conformational dynamics in a truncated epidermal growth factor receptor ectodomain. Biochemistry. 2011;50(23):5130–9.

55. Goldstein NI, Prewett M, Zuklys K, Rockwell P, Mendelsohn J. Biological efficacy of a chimeric antibody to the epidermal growth factor receptor in a human tumor xenograft model. Clinical cancer research: an official journal of the American Association for Cancer Research. 1995;1(11):1311–8.

56. Baselga J. The EGFR as a target for anticancer therapy—focus on cetuximab. European journal of cancer. 2001;37:16–22.

57. Mohan N, Luo X, Shen Y, Olson Z, Agrawal A, Endo Y, et al. A novel bispecific antibody targeting EGFR and VEGFR2 is effective against triple negative breast cancer via multiple mechanisms of action. Cancers. 2021;13(5):1027.

58. Makabe K, Yokoyama T, Uehara S, Uchikubo-Kamo T, Shirouzu M, Kimura K, et al. Anti-EGFR antibody 528 binds to domain III of EGFR at a site shifted from the cetuximab epitope. Scientific reports. 2021;11(1):1–6.

59. Shankaran V, Obel J, Benson III AB. Predicting response to EGFR inhibitors in metastatic colorectal cancer: current practice and future directions. The oncologist. 2010;15(2):157–67.

60. Simon N, Antignani A, Sarnovsky R, Hewitt SM, FitzGerald D. Targeting a cancer-specific epitope of the epidermal growth factor receptor in triple-negative breast cancer. JNCI: Journal of the National Cancer Institute. 2016;108(8).

61. Navabi H, Jasani B, Reece A, Clayton A, Tabi Z, Donninger C, et al. A clinical grade poly I:C-analogue (Ampligen) promotes optimal DC maturation and Th1-type T cell responses of healthy donors and cancer patients in vitro. Vaccine. 2009;27(1):107–15.

62. Salem ML, Kadima AN, Cole DJ, Gillanders WE. Defining the antigen-specific T-cell response to vaccination and poly(I:C)/TLR3 signaling: evidence of enhanced primary and memory CD8 T-cell responses and antitumor immunity. J Immunother. 2005;28(3):220–8.

63. Kershaw MH, Wang G, Westwood JA, Pachynski RK, Tiffany HL, Marincola FM, et al. Redirecting migration of T cells to chemokine secreted from tumors by genetic modification with CXCR2. Human gene therapy. 2002;13(16):1971–80.

64. Huang H, Xiang J. Synergistic effect of lymphotactin and interferon gamma-inducible protein-10 transgene expression in T-cell localization and adoptive T-cell therapy of tumors. International journal of cancer Journal international du cancer. 2004;109(6):817–25.

65. Kruse N, Moriabadi NF, Toyka KV, Rieckmann P. Characterization of early immunological responses in primary cultures of differentially activated human peripheral mononuclear cells. Journal of immunological methods. 2001;247(1-2):131–9.

66. Moser B, Wolf M, Walz A, Loetscher P. Chemokines: multiple levels of leukocyte migration control. Trends in immunology. 2004;25(2):75–84.

67. Alberts DS, Marth C, Alvarez RD, Johnson G, Bidzinski M, Kardatzke DR, et al. Randomized phase 3 trial of interferon gamma-1b plus standard carboplatin/paclitaxel versus carboplatin/paclitaxel alone for first-line treatment of advanced ovarian and primary peritoneal carcinomas: results from a prospectively designed analysis of progression-free survival. Gynecologic oncology. 2008;109(2):174–81.

68. Daniel D, Wilson NS. Tumor necrosis factor: renaissance as a cancer therapeutic? Current cancer drug targets. 2008;8(2):124–31.

69. Clark WH, Jr., Elder DE, Guerry Dt, Braitman LE, Trock BJ, Schultz D, et al. Model predicting survival in stage I melanoma based on tumor progression. Journal of the National Cancer Institute. 1989;81(24):1893–904.

70. Clemente CG, Mihm MC, Jr., Bufalino R, Zurrida S, Collini P, Cascinelli N. Prognostic value of tumor infiltrating lymphocytes in the vertical growth phase of primary cutaneous melanoma. Cancer. 1996;77(7):1303–10.

71. Amado RG, Wolf M, Peeters M, Van Cutsem E, Siena S, Freeman DJ, et al. Wild-Type KRAS is required for panitumumab efficacy in patients with metastatic colorectal cancer. Journal of Clinical Oncology. 2023;41(18):3278–86.

72. Ciardiello F, Tortora G. EGFR antagonists in cancer treatment. New England Journal of Medicine. 2008;358(11):1160–74.

73. Jonker DJ, O’Callaghan CJ, Karapetis CS, Zalcberg JR, Tu D, Au H-J, et al. Cetuximab for the treatment of colorectal cancer. New England Journal of Medicine. 2007;357(20):2040–8.

74. Randall KL. Rituximab in autoimmune diseases. Australian prescriber. 2016;39(4):131.

75. Zoeller JJ, Vagodny A, Daniels VW, Taneja K, Tan BY, DeRose YS, et al. Navitoclax enhances the effectiveness of EGFR-targeted antibody-drug conjugates in PDX models of EGFR-expressing triple-negative breast cancer. Breast Cancer Research. 2020;22(1):1–13.

76. Birkman E-M, Ålgars A, Lintunen M, Ristamäki R, Sundström J, Carpén O. EGFR gene amplification is relatively common and associates with outcome in intestinal adenocarcinoma of the stomach, gastro-oesophageal junction and distal oesophagus. BMC cancer. 2016;16(1):1–14.

77. Rakha EA, Tan DS, Foulkes WD, Ellis IO, Tutt A, Nielsen TO, et al. Are triple-negative tumours and basal-like breast cancer synonymous? Breast cancer research. 2007;9(6):1–3.

78. De Waele J, Verhezen T, van der Heijden S, Berneman ZN, Peeters M, Lardon F, et al. A systematic review on poly (I: C) and poly-ICLC in glioblastoma: adjuvants coordinating the unlocking of immunotherapy. Journal of Experimental & Clinical Cancer Research. 2021;40(1):1–20.

79. Edinger N, Lebendiker M, Klein S, Zigler M, Langut Y, Levitzki A. Targeting polyIC to EGFR over-expressing cells using a dsRNA binding protein domain tethered to EGF. PloS one. 2016;11(9):e0162321.

80. Sultan H, Wu J, Fesenkova VI, Fan AE, Addis D, Salazar AM, et al. Poly-IC enhances the effectiveness of cancer immunotherapy by promoting T cell tumor infiltration. Journal for immunotherapy of cancer. 2020;8(2).

81. Ogris M, Brunner S, Schüller S, Kircheis R, Wagner E. PEGylated DNA/transferrin– PEI complexes: reduced interaction with blood components, extended circulation in blood and potential for systemic gene delivery. Gene therapy. 1999;6(4):595–605.

82. Ogris M, Steinlein P, Kursa M, Mechtler K, Kircheis R, Wagner E. The size of DNA/transferrin-PEI complexes is an important factor for gene expression in cultured cells. Gene therapy. 1998;5(10):1425–33.

83. Garrett TP, McKern NM, Lou M, Elleman TC, Adams TE, Lovrecz GO, et al. Crystal structure of a truncated epidermal growth factor receptor extracellular domain bound to transforming growth factor α. Cell. 2002;110(6):763–73.

84. Abourbeh G, Shir A, Mishani E, Ogris M, Rodl W, Wagner E, et al. PolyIC GE11 polyplex inhibits EGFR-overexpressing tumors. IUBMB life. 2012;64(4):324–30.

85. Tolmachev V, Friedman M, Sandstrom M, Eriksson TL, Rosik D, Hodik M, et al. Affibody molecules for epidermal growth factor receptor targeting in vivo: aspects of dimerization and labeling chemistry. Journal of nuclear medicine : official publication, Society of Nuclear Medicine. 2009;50(2):274–83.

86. Nordberg E, Ekerljung L, Sahlberg SH, Carlsson J, Lennartsson J, Glimelius B. Effects of an EGFR-binding affibody molecule on intracellular signaling pathways. International journal of oncology. 2010;36(4):967–72.

